# Re-configuration of Chromatin Structure During the Mitosis-G1 Phase Transition

**DOI:** 10.1101/604355

**Authors:** Haoyue Zhang, Daniel J. Emerson, Thomas G. Gilgenast, Katelyn R. Titus, Yemin Lan, Peng Huang, Di Zhang, Hongxin Wang, Cheryl A. Keller, Belinda Giardine, Ross C. Hardison, Jennifer E Phillips-Cremins, Gerd A. Blobel

## Abstract

Higher-order chromatin organization such as A/B compartments, TADs and chromatin loops are temporarily disrupted during mitosis. These structures are thought to organize aspects of gene regulation, and thus it is important to understand how they are re-established after mitosis. We examined the dynamics of chromosome reorganization by Hi-C at defined time points following exit from mitosis in highly purified, synchronous cell populations. We observed that A/B compartments are rapidly established and progressively gain in strength following mitotic exit. Contact domain formation occurs from the “bottom-up” with smaller sub-TADs forming initially, followed by convergence into multi-domain TAD structures. CTCF is strongly retained at a significant fraction of sites on mitotic chromosomes and immediately resumes full binding at ana/telophase, the earliest tested time point. In contrast, cohesin is completely evicted from mitotic chromosomes and resumes focal binding with delayed kinetics. The formation of CTCF/cohesin co-anchored structural loops follows the kinetics of cohesin positioning. Stripe-shaped contacts anchored by CTCF grow in length, consistent with a loop extrusion process after mitosis. Interactions between cis-regulatory elements can form rapidly, preceding CTCF/cohesin anchored structural loops. Strikingly, we identified a group of rapidly emerging transient contacts between cis-regulatory elements in ana/telophase, that are dissolved upon G1 entry, co-incident with the establishment of inner boundaries or nearby interfering loops. Our findings indicate that distinct but mutually influential forces drive post-mitotic chromatin re-configuration to shape compartments, contact domains, cis-element contacts, and CTCF/cohesin dependent loops.

## Main text

The global restructuring of chromosomal architecture during the progression from mitosis into G1 phase provides an opportunity to examine hierarchies and mechanisms of chromosome organization ^1,2,3^. We performed *in situ* Hi-C experiments ^4^ at defined time points after mitosis following nocodazole induced pro-metaphase arrest-release in G1E-ER4 cells, a well-characterized subline of the murine erythroblast line G1E ^5^. To obtain synchronous cell populations we stably introduced into these cells a chimeric reporter consisting of mCherry fused to the murine cyclin B mitotic degradation domain (mCherry-MD), which is rapidly destroyed at the metaphase-anaphase transition (Fig. 1a, Extended Data Fig. 2a) ^6^. Fluorescence activated cell sorting (FACS) gated on mCherry and DNA content enabled isolation of highly homogeneous populations at ana/telophase (between anaphase and cytokinesis, ∼25min), early-(60min), mid-(120min) and late-G1 phase (240min after nocodazole removal) (Fig. 1a). To isolate pro-metaphase populations devoid of G2 cells, we employed a mitosis specific antibody (pMPM2) and reached a mitotic index of >98% as assessed by DAPI and lamin B1 staining (Extended Data Fig. 2b).

**Figure 1.**
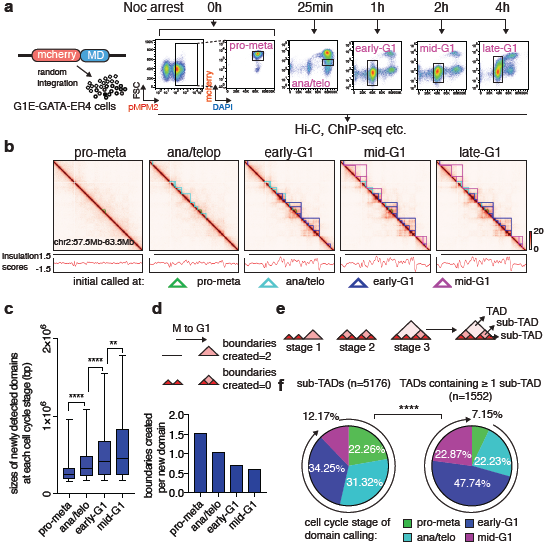
Contact domains develop from the “bottom-up” during the pro-M-G1 transition. **a**, Experimental setup. Left: the chimeric reporter gene encodes mcherry fluorescent protein and mouse cyclin B mitotic degradation domain. Right: flow chart showing the combination of nocodazole based synchronization-release strategy and fluorescence activated cell sorting (FACS) based on pMPM2 (pro-metaphase), or mcherry signal and DNA content. Boxes demarcate cell populations selected at each time point. **b**, Knight-Ruiz balanced, quantile normalized Hi-C interaction maps, coupled with insulation score track files (chr2:57.5M-63.5M). Domains emerging at each cell cycle stage are demarcated by color coded lines. Bin size: 10kb. **c**, size distributions of domains that are newly called at each cell cycle stage. 5-95% confidence intervals are shown. ** *p*<0.01, **** *p*<0.0001, Mann-Whitney test. **d**, Upper panel: two hypothetical cases of boundary creation with one extreme creating two boundaries per domain and another creating 0. Lower panel: ratio of the number of new boundaries and the number of new domains at each cell cycle stage. **e**, Schematic of hierarchically parsing contact domains into TADs or sub-TADs. **f**, Pie charts of the cell cycle distribution of sub-TADs (left) and TADs that contain *⩾* 1 sub-TADs based on their time of emergence. **** *p*<0.0001, Fisher’s exact test (pro-meta + ana/telo vs. early-G1 + mid-G1).

**Figure 2.**
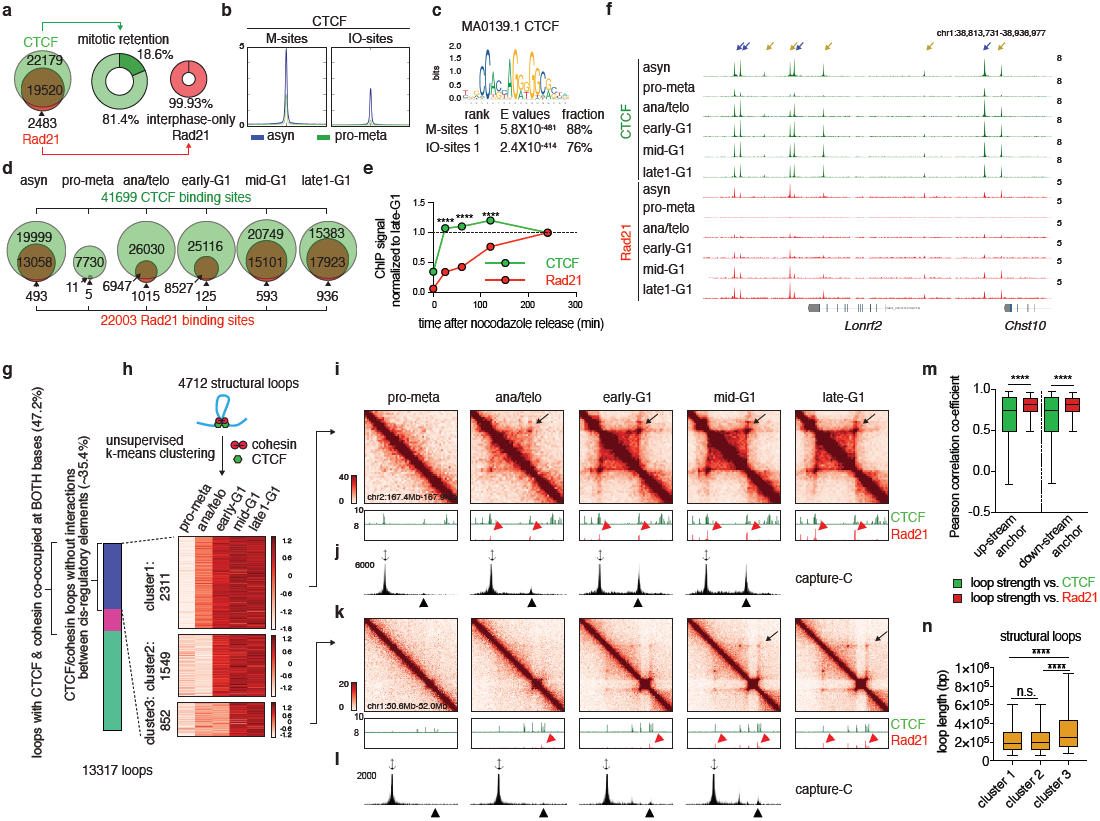
Focal accumulation of cohesin is delayed compared to CTCF and coincides with structural loop formation. **a**, Left: Venn diagram of the distribution of CTCF (green) and Rad21 (red) peaks. Middle & right: donut charts showing the percentages of mitotically retained CTCF (green) and Rad21 (red) peaks, respectively. **b**, Metagene plots of asynchronous and pro-metaphase CTCF ChIP-seq signals centered around mitotically retained (M) and interphase-only (IO) CTCF binding sites. **c**, Motif enrichment analysis of M and IO CTCF binding sites with indicated *E* values. **d**, Venn diagrams showing CTCF and Rad21 binding sites in asynchronous cells and across cell cycle stages. Pro-metaphase CTCF binding sites are defined as long as peaks are called in at least two biological replicates. Pro-metaphase Rad21 binding sites were included only when peaks were called in both biological replicates. **e**, Mean recovery curve of CTCF and Rad21. Each time point is normalized to late-G1. Error bars denote SEM. **** *p*<0.0001. Mann-Whitney test. **f**, genome browser tracks of CTCF and Rad21 binding at the *Lonrf2* and *Chst10* loci in asynchronous cells and across cell cycle stages. Blue and yellow arrows indicate M and IO CTCF binding sites, respectively. **g**, Bar graph showing the fraction of loops that belong to each indicated category. **h**, heatmap showing k-means clusters of the 4712 structural loops. **i**, KR balanced, quantile normalized Hi-C interaction maps showing a representative region that contains a cluster 1 structural loop (chr2: 167.4Mb-167.9Mb, loop indicated by black arrows) across all cell cycle stages, along with genome browser tracks showing CTCF and Rad21. Rad21 peaks that correspond to two bases of the loop are indicated by red arrowheads. Bin size: 10kb. **j**, Capture-C interaction profile of the same region as shown in (**i**). Anchor symbol shows position of the capture probe near the upstream loop anchor. Black arrowheads denote contacts. **k-l**, similar to (i-j) showing a representative region that contains a cluster 3 (slowly emerging) structural loop (chr1:50.6Mb-52.0Mb). **m**, Boxplots of Pearson correlation coefficients of structural loop strength and CTCF or Rad21 signal at up-stream or down-stream anchors. 5 to 95 percentile are plotted. **** *p*< 0.0001, Mann-Whitney test. **n**, Boxplots showing the length of cluster1, 2 and 3 structural loops. 5 to 95 percentile are plotted. **** *p*< 0.0001, Mann-Whitney test.

*In situ* Hi-C yielded ∼2 billion uniquely mapped interactions, with high concordance between biological replicates (Extended Data Fig. 2c-e). Consistent with previous studies, in pro-metaphase compartments and TADs are largely eliminated (Fig. 1b, Extended Data Fig. 3a) ^2,3,7^. In ana/telophase the earliest examined interval, transcriptionally active A and inactive B compartments are already detectable visually and by eigenvector decomposition (Extended Data Fig. 3a, b), and gain in intensity as cells advance into G1 (Extended Data Fig. 3c, d). These results are consistent with a previous report of early establishment of compartments after mitosis, using multiplexed 4C-seq ^8^. As expected, the A-type compartment is associated with active histone marks (Extended Data Fig. 3e) ^9,10^. As cells proceed towards late-G1, the characteristic checkerboard pattern of compartments visually expands away from the diagonal (Extended Data Fig. 3a). This feature is also seen in the progressively increased interactions between distally (>100Mb) separated genomic regions (Extended Data Fig. 3g). We quantified levels of compartmentalization at different distance scales across all cell cycle stages (methods) and found that ana/telophase cells display significantly reduced compartmentalization between genomic regions separated by >12Mb (Extended Data Fig. 3h). However, higher levels of compartmentalization are observed between more distal (>100Mb) genomic regions after G1 entry (Extended Data Fig. 3h, i), confirming that compartmentalization expands throughout the entire chromosome after mitosis. In summary, a major re-configuration of genome structure occurs during the pro-M-G1 transition, with a rapid establishment, progressive strengthening, and expansion of A/B compartments throughout the genome (Extended Data Fig. 1a).

**Figure 3.**
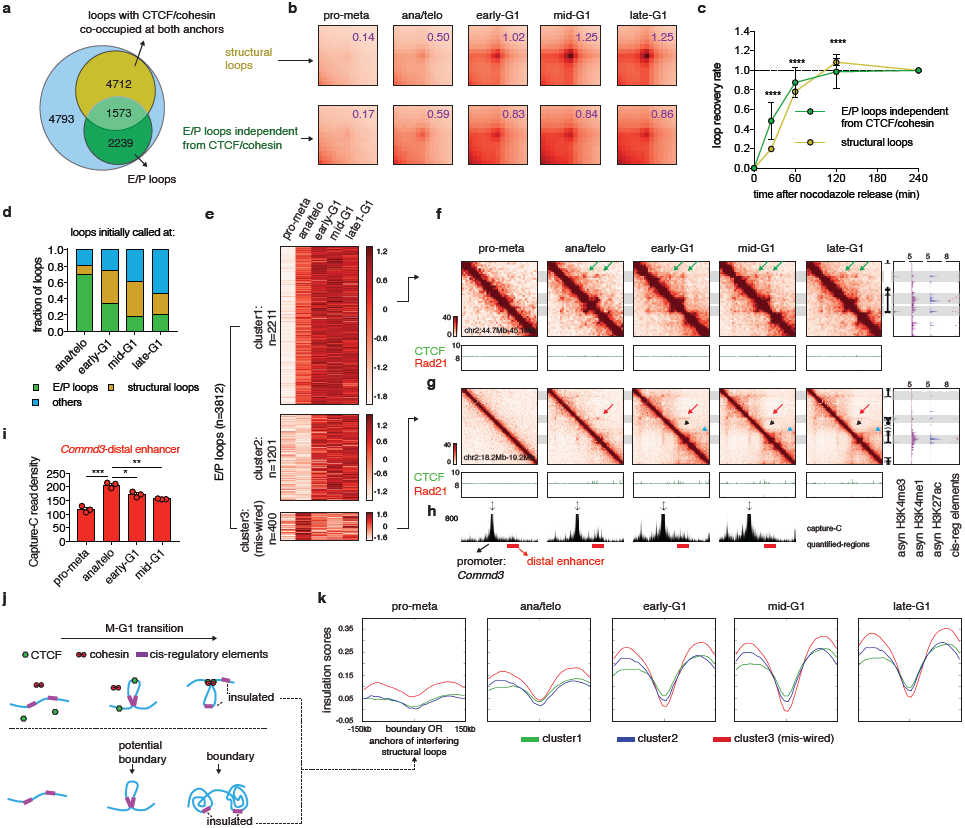
cis-regulatory contacts are established faster than structural loops and can be transiently mis-wired after mitosis. **a**, Venn-diagram of loops classified by CTCF/cohesin co-occupancy at both bases and enhancer-promoter (E/P) loops defined as the union of E-E, E-P and P-P contacts. **b**, Aggregated peak analysis (APA) from KR balanced, quantile normalized interaction maps of structural loops, and E/P loops that are independent of CTCF/cohesin. Numbers indicate average loop strengths (obs/exp) at each time point. Bin size: 10kb. **c**, Averaged recovery curves of structural loops and E/P loops independent of CTCF/cohesin. **** *p*<0.0001. Mann-Whitney test. **d**, Bar graphs showing the fraction of E/P loops, structural loops and “other” loops called at each cell cycle stage. **e**. Heatmap showing the results of k-means clustering on E/P loops. **f**, KR balanced, quantile normalized interaction maps of a representative region (chr2:44.7Mb-45.1Mb) containing a cluster 1 E/P loops (green arrows) across all cell cycle stages, coupled with tracks of CTCF and Rad21 occupancy. Tracks of H3K4me3, H3K4me1, H3K27ac, and annotations of cis-regulatory elements were from asynchronous cells. Bin size: 10kb. **g**, Representative region (*Commd3* locus, chr2:18.2Mb-19.2Mb) containing a cluster 3 (transiently mis-wired, red arrow) E/P loop. Black arrowhead indicates an emerging boundary between this mis-wired contact. Blue arrowhead indicates the formation of a potentially interfering structural loop. **h**, Capture-C interaction profiles of (**g**). Capture probe (anchor symbol) is near the *Commd3* promoter. Red bars demarcate the interaction between the *Commd3* promoter and down-stream enhancers. **i**, Bar graphs of Capture-C read densities of the red regions from (**h**). Error bars denote SEM from three biological replicates. * *p*<0.05, ** *p*<0.01, *** *p*<0.001 Student’s t test. **j**, Two potential mechanisms underlying transiently mis-wired E/P loops. **k**, Insulation profiles centered around the boundaries and interfering structural loop anchors that reside within cluster 1, 2 or 3 (mis-wired) E/P loops.

Next, we examined formation after mitosis of TADs and nested sub-TADs ^11,12^ using 3DNetMod ^13^. We identified 8073 contact domains progressively gained from pro-metaphase to mid-G1 (Fig. 1b, Extended Data Fig. 4b, Extended Data Table3). We observed gradually increased insulation at domain boundaries, suggesting strengthening of TADs over time (Extended Data Fig. 4a). Previous studies reported complete loss of domains in pro-metaphase ^2,3,7^. However, notably, despite significant attenuation of signal, residual domain-like structures are still qualitatively and algorithmically detectable in pro-metaphase cells (Extended Data Fig. 4C). The presence of pro-metaphase TAD/subTADs is unlikely due to contamination from interphase cells given the high purity of our mitotic preps and the fact that these residual structures are not homogenously distributed across the genome (Extended Data Fig. 4C). Thus, pro-metaphase is not prohibitive to the maintenance of chromatin structures.

Formation of TADs may occur via convergence of previously emerged sub-TADs (bottom-up), the partitioning of initially formed TADs into sub-TADs (top-down), or simultaneous birth of both contact domain types (Extended Data Fig. 4d). On average, contact domains called at time points later in G1 are larger than those called at preceding cell stages. (Fig. 1b, c). In addition, an average of ∼1.5 boundaries per domain are found at pro-metaphase, whereas only ∼0.58 boundary is created per domain in mid-G1 (Fig. 1d), suggesting that boundaries are being shared between sequentially formed domains. These results support the bottom-up model of domain reformation. To further test this model, we categorized all contact domains into 2897 TADs and 5176 sub-TADs, based on their topological hierarchy (Fig. 1e, methods). Among the former, 1552 contain at least one sub-TAD. Notably, higher proportions of sub-TADs are detected in pro-metaphase and ana/telophase compared to TADs that encompass them, suggesting that sub-TADs tend to assemble more rapidly (Fig. 1f). We next assessed the behavior of sub-TADs and TADs after their emergence. Once established, the majority of TADs remain unchanged without adding sub-divisions, arguing against the “top-down” model (Extended Data Fig. 4e). In contrast 85.4% and 69.2% of sub-TADs called in pro-metaphase and ana/telophase respectively, converge into larger domains during later stages (Extended Data Fig. 4f). Consistent with sub-TAD merging, we observed weakening of insulation at sub-TAD-boundaries (Extended Data Fig. 4g). To test whether this observation holds true across the genome, we computed the insulation scores of 1213 TAD-and 1139 sub-TAD-boundaries detected at pro-metaphase. For most TAD-boundaries insulation scores progressively decrease (signifying increased insulation) ^14^, whereas a significant portion of sub-TAD-boundaries display gradual increase of insulation scores (decreased insulation) (Extended Data Fig. 4h), indicating that flanking domains converge over time. Together, these analyses support a “bottom-up” model of hierarchical domain re-organization during the pro-M-G1 transition (Extended Data Fig. 1b).

A loop extrusion model has been proposed to explain the formation of TADs and chromatin loops, wherein the cohesin complex extrudes the chromatid until encountering a convergently oriented CTCF bound site ^15,16,17,18,19^. Since cell cycle dynamics of loop formation, CTCF and cohesin binding could inform this (or alternative) models, we surveyed the chromatin binding profiles of CTCF and cohesin by ChIP-seq. We generated highly concordant replicates (Extended Data Fig. 2f, g) and collectively identified 41699 CTCF and 22003 Rad21 (a cohesin subunit) binding sites (Fig. 2a, Extended Data Table. 6). ∼88.7% (19520) of Rad21 peaks are co-occupied by CTCF (Fig. 2a). Interestingly, ∼18.6% (7741) of CTCF peaks are reproducibly detected in pro-metaphase cells, suggesting significant amounts of mitotic retention (Fig. 2a, b, f, Extended Data Fig. 5a). Prior reports have described varying degrees of CTCF mitotic retention ^20,21,22^. This observation was confirmed by ChIP-qPCR at individual CTCF binding sites (Extended Data Fig. 5c, d). Motif enrichment analysis identified the canonical CTCF motif in both mitotically retained (M) and interphase only (IO) CTCF peaks (Fig. 2c). Unlike CTCF, Rad21 failed to show localized chromatin binding during pro-metaphase (Fig. 2a, f, Extended Data Fig. 5b-d). CTCF and cohesin resume chromatin occupancy after mitosis with remarkably different kinetics. The majority of CTCF peaks are restored in ana/telophase, whereas Rad21 peaks become detectable much more gradually (Fig. 2d-f, Extended Data Fig. 5e, f). This difference was confirmed by ChIP-qPCR where the signal was normalized to “input” chromatin (Extended Data Fig. 5g), and is consistent with a recent study using live cell imaging based approaches ^23^. Together, our results reveal distinct dynamics between CTCF and cohesin emergence. Notably, since ChIP-seq can only reliably detect focally enriched cohesin, our results do not permit inferences about the total amounts of cohesin on post mitotic chromatin that might not (yet) be focally localized. When viewed under this light, the results are compatible with parts of the loop extrusion model which predicates that CTCF halts cohesin movement.

The transient decoupling of cohesin from CTCF during mitotic exit offers the opportunity to separately assess their roles in post-mitotic loop formation. Using a modified HICCUP approach, we identified 13317 loops, progressively gained from pro-metaphase to late-G1, with a mean size of ∼213kb (Extended Data Fig. 6a, b, Extended Data Table 4). 6285 (∼47.2%) loops are anchored by CTCF and cohesin co-occupied binding sites at both bases. These loops were further filtered to eliminate interactions between putative cis-regulatory elements (i.e. enhancer – promoter loops), resulting in 4712 (∼35.4%) operationally defined “structural” loops (Fig. 2g, methods). To investigate how fast structural loops are formed, we performed k-means clustering, which revealed three clusters with distinct re-formation dynamics, consistent with the notion of highly variable recovery rates among loops (Fig. 2h) ^16^. Cluster 1 loops display strong interactions in ana/telophase, while formation of cluster 2 and 3 loops is delayed (Fig. 2h, i, k, Extended Data Fig. 6c-e). Capture-C ^24^ validated the differential dynamics of structural loops at two representative loci (Fig. 2j, l). Importantly, anchors of cluster 1 loops display early enrichment of Rad21 at ana/telophase, while anchors of cluster 2 and 3 loops acquire Rad21 more gradually (Fig. 2i, k, Extended Data Fig. 6d, e). In contrast, CTCF is enriched rapidly at anchors of all clusters of loops (Fig. 2i, k. Extended Data Fig. 6d, e). The strength of structural loops is highly correlated with Rad21 signal over time, but significantly less so with CTCF (Fig. 2m). Interestingly, cluster 3 loops also display significantly bigger loop sizes, suggesting that larger loops take longer to form (Fig. 2n). Together, these results reveal three clusters of structural loops with distinct re-formation dynamics. Our data also support a model that accumulation of cohesin, but not CTCF is limiting for structural loop formation after mitosis (Extended Data Fig. 1c).

Stripes in the contact maps are thought to reflect interactions between a single locus and a continuum of genomic regions and are considered as evidence of the loop extrusion model ^25^. We detected stripes using a modified version of the statistical modeling approach reported by Aiden and colleagues (methods)^25^. We identified 1,775 stripes genome-wide, the majority of which harbor inwardly oriented CTCF sites at their anchors (Extended Data Fig. 7a). Remarkably, these striped contacts grow directionally over time but display punctuated enrichment at selected CTCF sites (Extended Data Fig. 7b, d). This is consistent with an extrusion mechanism in which some CTCF bound sites serve as obstacles to further loop formation. We also observed blockage of stripe extension when it reaches to a group of strong CTCF binding sites, resulting in gradual formation of structural loops at the far end of the stripe (Extended Data Fig. 7b). Together, our data depict a picture of dynamic loop extrusion after mitosis (Extended Data Fig. 1d). Stripe like patterns that are established rapidly with little or no further growth were also observed and are discussed below (Extended Data Fig.7c, e).

Finally, we set out to investigate the re-formation of interactions between cis-regulatory elements. We identified 3812 chromatin loops, with both bases marked by promoters or putative enhancers, which we term as E/P loops. Note that this is probably an underestimate due to the limited resolution of Hi-C. Interestingly, a significant portion (∼58.7%, 2239) of E/P loops do not simultaneously harbor CTCF/cohesin co-occupied sites at both anchors, suggesting that they form independently of CTCF/cohesin (Fig. 3a). These seemingly CTCF/cohesin-independent E/P loops display stronger interactions in ana/telophase and significantly faster formation compared to CTCF/cohesin-anchored structural loops (Fig 3b, c). Accordingly, among loops detected in ana/telophase, ∼69.3% are E/P loops, while only ∼11.6% are structural loops (Fig. 3d). These two numbers are reversed to ∼18.4% and ∼42.3% in mid-G1, suggesting a trend of early establishment of E/P loops prior to structural loop formation after mitosis (Fig. 3d).

Clustering E/P loops according to their time of appearance yielded at least three classes with distinct post-mitotic formation kinetics. Cluster 1 (2211, ∼58%) E/P loops are rapidly enriched in ana/telophase, whereas cluster 2 (1201, ∼31.5%) assembles later in early-G1 (Fig. 3e, f, Extended Data Fig. 6f-h). Strikingly, we discovered a third cluster (400, ∼10.5%) of E/P loops that peak early in ana/telophase and gradually disappear in G1 (Fig. 3e, g, Extended Data Fig. 6h). We independently validated the transient nature of cluster 3 E/P loops by Capture-C at the *Commd3*, and two visually identified loci *(Pde12*, and *Morc3*) (Fig. 3h, i, Extended Data Fig. 6g, i). The transient nature of cluster 3 E/P loops led us to surmise that they represent inappropriate contacts. Interestingly, ∼55% of these transient E/P loops span either a future domain boundary and/or one anchor of a nearby interfering structural loop, which is why we speculate that they might be transiently “mis-wired” contacts (Fig. 3g, j, Extended Data Fig. 6g). Indeed, we observed pronounced strengthening of insulation at these boundaries or interfering loop anchors concurrently with the dissolution of the E/P loops (Fig. 3k). Conversely, interfering boundaries or loop anchors that reside in the cluster 1 and 2 E/P loops failed to display such progressive insulation (Fig. 3k). In sum, our data indicate that E/P loops can be mediated through mechanisms independent from CTCF and cohesin and can form faster than structural loops after mitosis (Extended Data Fig. 1e). We also identified a special class of transiently “mis-wired” E/P loops that are lost presumably due to subsequent insulation by CTCF and cohesin (Extended Data Fig. 1f).

We exploited the natural transition from a relatively unorganized state (pro-metaphase) into fully established chromatin organization late in G1 to interrogate mechanisms by which chromatin is hierarchically organized (Extended Data Fig. 1). We show that local (∼10Mb) compartmentalization of chromatin initiates rapidly after mitosis and continues to expand and increase in strength. Studying cell cycle dynamics of chromatin also enabled the testing of predictions made by the loop extrusion model. First, larger TADs and structural loops are completed more slowly than smaller ones. Second, the formation rate of pairwise interactions between CTCF and cohesin binding sites decreases with distance between them (Extended Data Fig. 8a). Third, chromatin stripes increase in length over time. Forth, based on the kinetics of CTCF and cohesin deposition on chromatin, it is clear that CTCF does not form detectable loops without cohesin even though it can multimerize ^26^. However, it is possible that CTCF pairs with itself or other factors such as YY1 to establish contacts among cis regulatory elements such as those observed at early time points independently of cohesin ^27,28^.

Our integrative analysis of loops and histone modification profiles reveals a group of E/P loops that can be independent from CTCF and cohesin co-binding. Their most distinctive features are their fast appearance compared to structural loops. It is possible that these contacts form via collisions of chromatin regions with similar epigenetic states, which is supported by our observation that their post-mitotic recovery rate positively correlates with the intensity of active histone marks at anchors (Extended Data Fig. 8b, c). Intriguingly, 16.4% of stripe-like structures that lack inwardly oriented CTCF at their anchors display only little or no further growth during G1 phase and are highly enriched for H3K27ac at their anchors (Extended Data Fig. 7c, e, f). Loop extrusion unlikely accounts for these stripe shaped contacts and they might instead be mediated by rapid chromatin segregation, or facilitated by local enrichment of transcription factors or co-activators ^29^. Similarly, transiently mis-wired E/P loops might reflect less discriminatory affinity among regions with similar chromatin states. Hence, a transient cellular state during which cis-elements “forget” their proper chromatin contacts is overcome by the formation of boundaries or interfering structural loops. It is important to note that the transcriptional reactivation dynamics of a particular gene does not necessarily mirror that of the interaction between its promoter and distal enhancers. In fact, spiking of expression has been observed on genes that display more gradually increased promoter-enhancer interactions ^6^. In summary, our findings describe a dynamic hierarchical framework of post-mitotic chromatin configuration that supports a bottom-up model for the formation of contact domains, implicates CTCF and cohesin in post-mitotic loop extrusion, and identified extrusion independent pathways that lead to compartmentalization and contacts of cis-regulatory networks.

## Acknowledgements

We thank members of the Blobel and Phillips-Cremins labs for helpful discussions. We thank Effie Apostolou and Job Dekker for discussing data prior to publication. This work was supported by grants R37DK058044 to G.A.B.; R24DK106766 to G.A.B. and R.C.H.; U01HL129998A to J.E.P.C. and G.A.B.; the Alfred P. Sloan Foundation to J.E.P.C.; and a generous gift from the DiGaetano family to G.A.B. J.E.P.C. is a New York Stem Cell Foundation (NYSCF) Robertson Investigator. We thankfully acknowledge the support by the Spatial and Functional Genomics program at The Children’s Hospital of Philadelphia.

## Author Contributions

H.Z., J.E.P.-C., and G.A.B conceived the study and designed experiments. H.Z. performed experiments with help from P.H., H.W., C.A.K., B.G. and B.C.H. D.J.E. performed initial Hi-C data pre-processing and domain calling. H.Z. performed A/B compartment and ChIP-seq related analysis with help from Y.L. T.G.G performed loop calling and K.R.T performed stripe calling related analysis. D.Z. contributed to Capture-C related analysis. H.Z., J.E.P.-C., and G.A.B wrote the paper with input from all authors.

## METHODS

### Experimental procedure

#### Cell line, pro-metaphase synchronization and release and cell sorting

The erythroblast subline G1E-ER4 was cultured as previously described ^5^. Cells were transduced with retrovirus expressing mcherry-MD. Cells with positive mcherry signal was sorted as a pool through fluorescent activated cell sorting (FACS) and then further expanded. To synchronize cells at pro-metaphase, actively proliferating cells were treated with 200ng/ml nocodazole for 7-8.5h at a density of 0.6-0.8 million cells per ml. To release cells from mitosis, cells were washed once and then immediately re-suspended with warm nocodazole free culture medium for a variety of durations. Cells were released for 0min, 25min, 60min, 120min and 240min to enrich pro-meta, ana/telo, early-G1, mid-G1 and late-G1 populations respectively. Crosslinking and cell sorting were performed as previously described ^6^. Briefly, at the time of collection, cells were re-suspended in 1×PBS at a density of ∼50million/36ml and crosslinked with formaldehyde (1% for ChIP and 2% for *in-situ* HiC and capture-C) for 10min, followed by 5min quenching with 1M glycine. For *in-situ* HiC and capture-C experiments, fixed cells were re-suspended in permeabilization buffer (1×PBS, 2mM EDTA and 0.1% Triton X-100) for 5min and then re-suspended in 1×FACS sorting buffer (1×PBS, 2% FBS, 2mM EDTA and 0.02% NaN_3_) containing 20ng/ml DAPI. For ChIP-seq and ChIP-qPCR experiments, every 50million of fixed cells were re-suspended in 1ml of permeabilization buffer containing protease inhibitors (Sigma, P8340), 1mM phenylmethylsulfonyl fluoride (PMSF) and anti-pMPM2 (Millipore, 05-368) antibody with a concentration of 0.2 μl/10million cells. Cells were incubated at room temperature for 50min. Cells were then further stained with F(ab’)2-goat anti-mouse secondary antibody (Thermo Fisher Scientific, 17-4010-82) at a concentration of 2 μl/10million cells and incubated at room temperature in dark for 30min. Finally, cells were re-suspended in 1×FACS sorting buffer containing protease inhibitor, PMSF and 20ng/ml DAPI and subject to cell sorting. Cells were sorted with designated gates based on mcherry and DAPI signal (Fig. 1a) at different time points with a Beckman Coulter Moflo Astrios sorter. For pro-metaphase enrichment, 4N populations with pMPM2 positive staining and high mcherry signal were collected. For lamin B1 fluorescent staining and imaging, 1-2 million sorted cells were stained with anti-lamin B1 (Abcam, ab16048) antibody at a concentration of 0.5μl/million cells. Cells were spun onto slides with a cyto spinner, mounted and imaged with a Leica DM6000 motorized upright microscope.

#### *In-situ* Hi-C

3C libraries were performed as previously described ^4,30^. Briefly, 5million pro-meta or ana/telo cells and 10million interphase cells were collected from cell sorting per experiment. Cells were pelleted, re-suspended in 1ml cold Cell Lysis Buffer (10mM Tris pH 8, 10mM NaCl, 0.2% NP-40/Igepal) and incubated on ice for 10min. Cell nuclei was pelleted at 4°Cand washed with 800μl 1.2μDpnII buffer. Nuclei was pelleted at 4°Cand re-suspended in 500μl 1.2μDpnII buffer. 15.5μl SDS (10%) was added into the system to reach a final concentration of 0.3%. Samples were then incubated at °C on a thermomixer for 1h with shaking at 950rpm. 50μl Triton X-100 (20%) was added to the system to reach a final concentration of 1.8%. Samples were further incubated at °C on a thermomixer for 1h at 950rpm. 300U of DpnII restrictive enzyme (NEB, R0543M) was then added into the system and samples were incubated at 37°Cfor overnight with shaking at 950rpm. Another 300U of DpnII was added into the system and samples were further incubated at °C for 2-4h with shaking at 950rpm. Samples were then incubated at 65°C for 20min with shaking at 950rpm to inactivate DpnII. Nuclei was pelleted, cooled down to room temperature, re-suspended in 300μl 1×NEB buffer2, containing 50μM biotin-14-dATP (Thermal Fisher Scientific, 19524016), dGTP, dCTP, dTTP (Promega, U121B, U122B, U123B) and 40U DNA Polymerase I, Large (Klenow) fragment (NEB, M0210) and incubated at °C for 1.5h with shaking at 950rpm. The system was then added with 120μl 10×T4 DNA ligase buffer, 12μl 100×BSA, 758μl nuclease free water and 10μl (4000U) T4 DNA ligase (NEB, M0202M). Ligation was carried out at 16°C for 4h and room temperature for additional 2h. 20μl of 20mg/ml protease K (3115879 BMB) and 120μl of 10%SDS were added into the system and samples were de-crosslinked at 65°C for overnight. An additional 10μl of protease K was added and samples were further de-crosslinked at 55°C for 2h. DNase-free RNase was added into the system and samples were incubated at °C for 30min. Ligated DNA was extracted with phenol chloroform extraction and re-constituted in 180μl nuclease free water. DNA concentration was quantified with a Qubit 4 Fluorometer (Thermal Fisher Scientific, Q33226). Ligated DNA was sonicated to 200-300bp (Epishear, Active Motif, 100% amplitude, 30s ON and 30s OFF, 25-30min) and cleaned up with AMPure XP beads (Beckman Coulter). Each sample of biotinylated DNA was pulled down with 100μl Dynabeads MyOne Streptavidin C1 beads (Thermal Fisher Scientific, 65002) by incubating at room temperature for 15min. The amount of bound DNA was calculated by subtracting the amount of unbound DNA from the amount of total DNA. DNA bound streptavidin beads were washed once with 1×bead wash buffer (5mM Tris-HCl (pH 7.5), 0.5mM EDTA and 1M NaCl) and once with T4 DNA ligase buffer. Streptavidin beads were re-suspended with 84μl water. Library was prepared using NEBNext DNA Library Prep Master Mix Set for Illumina (NEB E6040, M0543L, E7335S) according to manufacture instruction. On-bead end repair was performed by mixing 84μl biotinylated beads with 10μl NEBNext End Repair Reaction Buffer (10×), 5μl NEBNext End Repair Enzyme Mix and 5U DNA Polymerase I, Large (Klenow) Fragment (NEB, M0210). The mixture was incubated at room temperature for 30min. Beads with end repaired blunt DNA was washed once with 600μl 1×bead wash buffer and then 500μl 1×NEB buffer2. Beads was re-suspended in 42μl nuclease free water. Beads were then mixed with 5μl NEBNext dA-tailing reaction buffe (10×) and 3μl Klenow Fragment (3’→5’ exo-) for dA-tailing. The mixture was incubated at °C for 30min. Beads were then washed once with 600μl 1×bead wash buffer and then 600μl 1×T4 DNA ligase buffer. Beads were re-suspended in 20μl nuclease free water. Adaptor ligation was performed by mixing the beads with 10μl Quick Ligation reaction buffer (5×), 15μl NEBNext Adaptor (15μM) and 5μl Quick T4 ligase. The mixture was incubated at room temperature for 15min. 3μl of USER enzyme was then added into the mixture and incubated at °C for 15min. Beads were washed twice with 600μl 1×bead wash buffer and re-suspended in 50μl nuclease free water containing 0.1% SDS. Beads were incubated at 98C for 10min to release DNA and DNA was cleaned up with AMPure XP beads. Purified adaptor ligated DNA was then index labeled with NEBNext multiplex oligos for 6 cycles on a thermal cycler. PCR products were purified using AMPure XP beads. Sequencing was performed on an Illumina NextSeq 500 platform using Illumina sequencing reagents according to manufacturer’s instructions.

#### Capture C

Capture-C experiments were performed as previously described ^24,30^. 3C library preparation was similar to that of *in-situ* HiC experiments without biotin nucleotide labeling. Briefly, DNA was directly ligated after DpnII digestion. Ligated DNA was purified, fragmented and converted into 3C library using NEBNext DNA Library Prep Master Mix Set for Illumina (NEB E6040, M0543L, E7335S). After indexation with NEBNext multiplex oligos, DNA was cleaned up with AMPure XP beads and further amplified with Illumina P5 and P7 primers for 6 cycles. To perform oligo pull-down, 3C libraries were completely dried on a thermal cycler at 60C. Dried DNA was re-constituted with 13μl hybridization mixture (2.5μl nuclease free water, 7.5μl 2×hybridization buffer, 3μl hybridization component A, Nimblegen SeqCap Hybridization and Wash kit) for 10min at room temperature. DNA was denatured at 95°C for 10min and then added with 2μl 3pmole biotin-labeled capture probe pool (probe sequence available in Extended Data Table 1) to make a 15μl hybridization system. Hybridization was immediately carried out at 47C for more than 24h on a thermal cycler. Hybridized DNA was pulled down with Dynabeads MyOne Streptavidin C1 beads (40μl per capture) and washed with Nimblegen SeqCap Hybridization and Wash kit according to manufacture instruction. Captured DNA was then eluted into 25μl 0.125N NaOH, followed by HCl neutralization. Captured DNA was then purified with AMPure XP beads. Purified DNA was then amplified with Illumina P5 and P7 primer for 12 cycles and PCR products were cleaned up with AMPure XP beads. PCR products were dried on a thermal cycler at 60°C to initiate a similar second round of capture for further enrichment of targeted DNA. DNA was eluted into 25μl nuclease-free water and subject to sequencing with Illumina NextSeq 500 platform using Illumina sequencing reagents according to manufacturer’s instructions.

#### ChIP-seq and ChIP-qPCR

Chromatin immunoprecipitation (ChIP) was performed with anti-CTCF (Millipore, 07-729) and Rad21 (Abcam, ab992) antibodies. Crosslinking of samples was performed in the sample preparation step for sorting. 7-10million cells per sample was collected from sorting, pelleted and stored at -80°C. Upon experiment, sorted samples were re-suspended in 1ml Cell Lysis Buffer (10mM Tris pH 8, 10mM NaCl, 0.2% NP-40/Igepal) containing fresh protease inhibitors (Sigma P8340) and 1mM phenylmethylsulfonyl fluoride (PMSF) for 20min on ice. Nuclei was pelleted at 4°C and re-suspended in 1ml Nuclear Lysis Buffer (50mM Tris pH 8, 10mM EDTA, 1% SDS) containing fresh protease inhibitors and PMSF and incubated for 20min on ice. Samples were added with 0.6ml IP Dilution Buffer (20mM Tris pH 8, 2mM EDTA, 150mM NaCl, 1% Triton X-100, 0.01% SDS) containing fresh protease inhibitor and PMSF and then subject to sonication for 45min (Epishear, Active Motif, 100% amplitude, 30s ON and 30s OFF, ∼45min).

After sonication, samples were spun at 21130g at 4°for 10min to pellet cell debris. Supernatant was transferred to a new tube and supplemented with 3.4ml IP Dilution Buffer containing fresh protease inhibitor, PMSF, protein A/G agarose beads (prepared by mixing protein A (Thermo Fisher Scientific 15918014) and protein G (Thermo Fisher Scientific 15920010) agarose beads at 1:1 ratio) and 50μg isotope matched IgG. Chromatin preclear was carried out for over 2h at 4°C. 200μl (4%) of precleared chromatin was taken out from subsequent IP steps and set as input. Pre-cleared chromatin was then added to protein A/C beads bound with antibodies and incubated overnight with rotation at 4°C. After IP, beads were washed once with IP Wash Buffer I (20mM Tris pH 8, 2mM EDTA, 50mMNaCl, 1% Triton X-100, 0.1% SDS), twice with High Salt Buffer (20mMTris pH 8, 2mM EDTA, 500mMNaCl, 1% Triton X-100, 0.01% SDS), once with IP Wash Buffer II (10mMTris pH 8, 1mM EDTA, 0.25 M LiCl, 1% NP-40/Igepal, 1% sodium deoxycholate) and twice with PE Buffer (10mM Tris pH 8, 1mM EDTA pH 8). All washing and pelleting steps were performed on ice or at 4°C. Beads were then moved to room temperature and eluted twice with 100μl freshly prepared Elution Buffer (100mM NaHCO3, 1%SDS) to reach a final concentration of 200μl. Eluted samples were supplemented with 12μl 5M NaCl, 2μl DNase-free RNaseA (10mg/ml, 10109169001 BMB) and incubated at 65°C for overnight. Samples were then added with 3μl protease K (20mg/ml, 3115879 BMB) and further de-crosslinked at 65°C for 2h. De-crosslinked DNA was added with 10μl 3M sodium acetate pH 5.2 and purified with QIAquick PCR Purification kit (QIAGEN 28106) according to manufacture instruction. Similar de-crosslinking and purification steps were performed on input samples. ChIP-qPCR was performed with Power SYBR green master mix (Thermal Fisher Scientific) on an ABI Vii7 real-time PCR machine. All primer sequences are listed in Extended Data Table 1. For ChIP-seq experiments, purified IP or input DNA was subject to library construction for Illumina sequencing. Library construction was performed using Illumina’s TruSeq ChIP sample preparation kit (Illumina, catalog no. IP-202-1012) according to manufacturer’s specifications with the addition of size selection (left side at 0.9x, right side at 0.6x) using SPRIselect beads (Beckman Coulter, catalog no. B23318) prior to PCR amplification. Library size was determined (average 351 bp, range 333-372 bp) using the Agilent Bioanalyzer 2100, followed by quantitation using real-time PCR using the KAPA Library Quant Kit for Illumina (KAPA Biosystems catalog no. KK4835). Libraries were then pooled and sequenced on the Illumina NextSeq 500 platform using Illumina sequencing reagents according to manufacturer’s instructions.

### Quantification and Data Analysis

#### Hi-C read alignment and matrix assembly

Paired-end reads were aligned independently to the mm9 mouse genome using bowtie2 (global parameters: --very-sensitive –L 30 –score-min L,-0.6,-0.2 –end-to-end --reorder; local parameters: --very-sensitive –L 20 –score-min L,-0.6,-0.2 –end-to-end --reorder) through the HiC-Pro software ^31^. Unmapped reads, non-uniquely mapped reads, and PCR duplicates were filtered and uniquely aligned reads were paired (Extended Data Table 2). We assembled raw contact matrices for all replicates and time points into 10 kb non-overlapping bins. The number of intra-chromosomal contacts for 10 kb matrices are provided in Extended Data Table 2.

#### Hi-C inter-replicate correlations

For each pair of replicates, we computed the Pearson’s correlation coefficient between entries of the 250 kb binned raw contact matrices. We excluded all *trans* interactions, as well as bin-bin pairs for which zero interactions were detected in either replicate. The pairwise Pearson’s correlation coefficients are shown in (Extended Data Fig. 2c). For each time point, we plotted a hexbin plot of all the *cis* 250 kb bin-bin pairs with non-zero raw read counts in both replicates, setting the y-coordinate of each bin-bin pair to its raw read count in replicate 1 and its *x*-coordinate to its raw read count in replicate 2 (Extended Data Fig. 2e). The high correlation between biological replicates across different conditions allowed us to merge their reads for down-stream analysis.

#### A/B compartment detection by eigenvector decomposition

“.hic” files of the merged reads (converted from validPair files through HiC-Pro) across all cell cycle stages were used as input for compartment calling. Compartments were quantified by eigenvector decomposition on the Pearson’s correlation matrix of the observed/expected value of 250kb binned, Knight-Ruiz balanced cis-interaction map (achieved through the Eigenvector utility of juicer_tools_1.6.2 ^32^). Compartment tracks for each individual chromosome were represented by the values of cis-eigenvector 1 (dominant principle component of the Pearson’s correlation matrix). The direction cis-eigenvector 1 was arbitrary. Positive and negative values were assigned to A-(active) and B-(inactive) compartments respectively based on their association with gene density. Cis-eigenvector 1 of each individual biological replicate at each cell cycle stage was also computed. Pearson correlation of cis-eigenvetor 1 values across all replicates and time points were plotted in Extended Data Fig. 2d.

#### Saddle plotting and compartment strength calculation

To calculate the strength of compartmentalization at each cell cycle stage, we firstly obtained the observed/expected values of each 250kb bin after Knight-Ruiz balancing from the merged .hic files through the Dump utility of juicer_tools (1.6.2) ^32^. We then, re-ordered each column and row of the observed/expected matrix of late-G1 phase based on the their cis-eigenvector 1 values, so that they were arranged in an ascending order from left to right and from top to bottom. In this way, Bins associated with B-type compartments were moved to the upper-left corner while those associated with A-type compartments were arranged to the lower-right corner. This order of bins from late-G1 phase was then applied to every other cell cycle stages. Bins across the entire chromosome were then aggregated into 50 equally sized sections to create the saddle plot. Compartment strength of each individual chromosome was calculated as the following: compartment strength = (median(20% strongest AA)+median(20% strongest BB))/(median(20% strongest AB)+median(20% strongest BA)).

#### Distance dependent contact probability curve *P(s)*

*P(s)* curve illustrates contact frequency *P* between two genomic loci as a function of their genomic distance *s*. To calculate *P(s)*, we employed the Knight-Ruiz balanced cis-contact matrix with 10kb binning. For each individual chromosome, we then determined a series of distances starting from 10kb that grew exponentially with a factor of 1.12, floored to the nearest integer. For each distance (10kb, 20kb, 30kb……), we calculated contact probability through dividing aggregated read counts within that distance by total read counts within that chromosome. Similar approach was performed for all chromosomes across all cell cycle stages as well as asynchronous sample ^33^. Chromosome averaged contact probability was used to represent genome-wide distance dependent contact probability. Contact probability curve of each cell cycle stage was smoothened through LOWESS fitting in R.

#### Quantification of A/B-compartment expansion: calculation of *R(s)*

Visual inspection of Hi-C contact maps suggests spreading of the plaid-like compartmentalization pattern over time. We introduced a new metric *R(s)* to quantify the level of compartmentalization at any genomic distances *s* (Extended Data Fig. 3f). *R(s)* was computed on the 250kb binned, Knight-Ruiz balanced observed/expected maps. For a 250kb interacting bin-bin pair separated by a defined genomic distance *s* (eg. 1Mb or 100Mb), we computed the product of the two cis-eigenvector 1 values that correspond to both bin of that pair. We then, calculated *R* as the Spearman correlation coefficient between cis-eigenvector 1 products and the observed/expected values of all bin-bin pairs sharing the same genomic separation chromosome wide (Extended Data Fig. 3f). *R* was calculated for a series of genomic separations (*s*=250kb, 500kb, 750kb … 125Mb) to generate the distance dependent *R(s)* curve for each individual chromosome. Same calculation was applied to all cell cycle stages as well as asynchronous sample^33^. In highly compartmentalized regions, interacting bin-bin pairs between the same type of compartments (A-to-A or B-to-B) tend to have high observed over expected values as well as positive (>0) cis-eigenvector 1 products, whereas interacting bin-pairs between different type of compartments (A-to-B or B-to-A) usually have low observed over expected values and negative (<0) cis-eigenvector 1 products. Thus, *R(s)* is expected to be high in well-compartmentalized regions. In pro-meta and ana/telo samples, interactions significantly diminish as *s* increases to diagonal distal regions, resulting in more randomized noisy observed/expected values. As a result, *R(s)* significantly drops when *s* increases beyond 10 to 20Mb. After G1 entry, *R(s)* becomes more persistent as s increases, suggesting that previously un-organized regions start to form the plaid like compartment pattern.

#### Domain calling preprocessing

For domain calling, we performed Knight-Ruiz matrix balancing on 10 kb matrices consisting of merged replicates for every condition and time point using Juicebox (juicer_tools_0.7.0) for each chromosome individually with no row removal. The one exception was chromosome 8, which required removal of sparse rows with less than 3 nonzero counts to achieve matrix balancing. We performed quantile normalization across all merged replicates to equalize global counts distributions across all timepoints while preserving the underlying biological differences unique to each time point. Prior to quantile normalization, if less than 50 percent of merged replicates have a nonzero count across timepoints for bin-bin interaction, the count is removed from consideration for all timepoints. Ties, in which counts have same starting value, were assigned the average value of the lowest rank in the set of tied counts.

#### Domain calling

We identified TADs and subTADs genome-wide using our recently published 3DNetMod method ^13^ (https://bitbucket.org/creminslab/3dnetmod_method_v3.0_development). The same parameters were implemented across all time points on quantile normalized, merged replicates at 10 kb resolution.

Genome-wide counts data for each merged replicate were log transformed and chunked into 7.5 Mb regions with 5 Mb overlap. We analyzed all regions except those that (1) overlapped with telomeres or centromeres, (2) exhibited consecutive zero counts on the diagonal for a genomic distance of >= 500 kb, or (3) exhibited zero counts for >= 1/3 of all pixels on diagonal. To select gamma, we chose a minimum plateau size of 30 (i.e. at least 30 consecutive gamma values, every 0.01 gamma step, with same mean number of called domains per 20 attempts, at each successive gamma). A sweep of gamma values was selected for domain detection as the mean gamma at every plateau. Domains were identified at each gamma by running 3DnetMod 20 times (i.e. 20 partitions) and computing a consensus via the adjusted rand index. After concatenating all unique, domains genome-wide, we filtered those that were smaller than 150 kb. Domains had to be established with boundary at least 4 bin distance from chunked region edge. We also filtered low-confidence domains by computing the variation of the domain partitions compared to the the partition block consensus. If domains exhibited a spatial variance of > 12 bins on both left and right boundaries, they were removed from consideration. The final list of domain calls were merged to account for redundant, nearly fully overlapping domains. Domains that overlapped with only a +/-1 bin fluctuation on both boundaries were merged into a single domain bounded by start and end coordinates separated by the largest genomic distance.

We next classified the domains by their unique emergence at each time point. To be classified as “time point emergent”, a domain uniquely emerging at a specific time point (0, 25, 60, and 120 minutes) also had to persist for at least one future time point. To determine if a domain existed at a later time point, we required that it either (1) overlapped the original domain with a +/- 6 bin fluctuation on both boundaries or (2) the difference in base pairs between the two left boundaries and the two right boundaries was less than 10% of the original domain size. The domain added to a specific “time point emergent” class exhibited the coordinates for the original emergent domain, not the coordinates of the later time point. Because 240 minutes has no future time point for comparison, emergent domains identified only at 240 minutes were dropped from consideration. Emergent domains identified at 0, 25, 60, and 120 minutes were listed in Extended Data Table 3.

To create final lists of domains with “time point emergent” shared unique boundaries, we then implemented a final step in which we adjusted the domain boundaries to ensure that domains with highly similar boundaries shared the same boundary. We compared left and right boundaries of each emergent domain separately to all other boundaries of emergent domains. If domains are accessed to share same boundary (i.e. boundary coordinates are close), we reassign the same boundary coordinate to the domains under consideration. More specifically, If the base pair distance between the matched, similar boundaries is less than 7.5% of domain size for all domains under consideration or within 3 bins, a final averaged unique boundary is reassigned to all emergent domains. Final unique, non-redundant boundaries emergent at 0, 25, 60, and 120 minutes were listed in Extended Data Table 3.

#### Insulation Score

A 120 kb square window (12×12 bins) with one bin offset from the diagonal was tiled across the genome on merged, matrix balanced Hi-C maps at each time point. Counts in the 12×12 bin window were summed, normalized by the chromosome-wide mean IS, log transformed, and recorded as the Insulation Score (IS). Windows spanning low information content regions (< 12 counts or IS window is interrupted by the end of the chromosome) were discarded from the analysis.

#### Domain and boundary partition

**TAD & sub-TAD.** All domains were categorized into either TADs or sub-TADs based on their topological hierarchy. Briefly, domains across all cell cycle stages were summarized into a union domain list irrespective of their time of detection (Fig. 1e). If a domain is completely encompassed by any other domains, it was classified as a sub-TAD. If a domain is not completely encompassed (no overlap or only partially overlap) by other domains, it was classified as a TAD. Boundaries may be associated with multiple domains simultaneously. All boundaries were partitioned into two groups: TAD boundaries and sub-TAD boundaries. TAD boundaries are boundaries not encompassed by any other domains or boundaries encompassed by other domains but only associated with TADs (no sub-TAD association). sub-TAD boundaries are boundaries completely encompassed by other domains and associated with at least one sub-TAD.

#### Loop and Stripe Preprocessing

For loop and stripe identification, we implemented a filter that removes poorly mapped regions based on the mm9 75-mer CRG Alignability track from ENCODE. Specifically, we discarded a 10 kb bin if the average mappability of a 50 kb window centered on that bin was below 50%. In addition to a mappability filter, we also implemented a high outlier filter to further prevent balancing artifacts. We removed pixels in the raw contact matrices that exhibit high fold changes relative to a neighborhood of adjacent pixels after balancing. The neighborhood around a given pixel was defined by a 5×5 square footprint, which was used to determine a local median. A given pixel was determined to be an outlier if its value was either greater than 4 or greater than 4 multiplied by the local median, whichever threshold was greater. After filtering, we performed Knight-Ruiz matrix balancing on 10 kb matrices of individual, non-merged replicates for every condition and timepoint using Juicebox (juicer_tools_0.7.0) for each chromosome individually with no row removal.

#### Stripe Identification

To call loop extrusion lines in 10 kb balanced matrices, we implemented a modified version of the stripe calling pipeline published previously ^25^. We first computed local horizontal and vertical signals from the raw, unbalanced contact matrix, *X*, by finding the median (*M*) of 10 pixels orthogonal to each query pixel, *x_i,j_* (**Equations 1-4**). For the vertical signals, the median was calculated with respect to 5 pixels directly above and below the primary ij pixel. For the horizontal signals, the median was calculated with respect to 5 pixels directly left and right of the primary ij pixel. We captured the signal using a median because it produced more precise estimations of local vertical and horizontal signals than the summation approach originally reported ^25^.

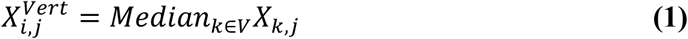

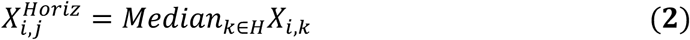

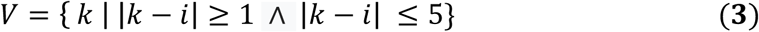

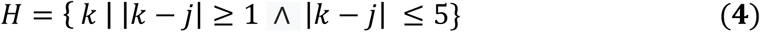

The background expected values for the vertical and horizontal signals were computed as described by Vian et al. ^25^. Briefly, we evaluated the local neighborhood around each pixel by computing the median counts value in the balanced contact matrix, *S*, using top, bottom, left, and right footprints (**Equations 5-12**).

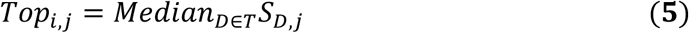

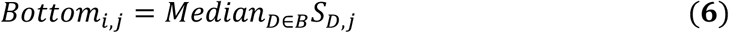

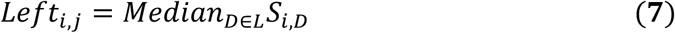

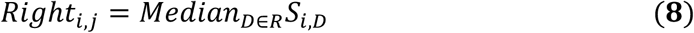

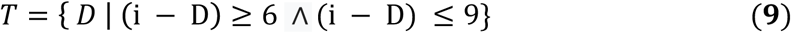

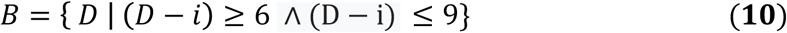

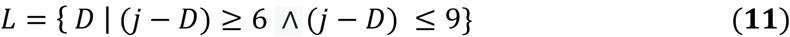

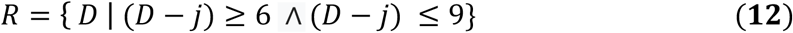

Next, we computed the vertical and horizontal signals on the balanced contact matrix, *S*, using the same logic described in **Equations 1-4** to ultimately compute the bias factors that allow deconvolution of the directional footprints to pseudo raw counts for statistical testing (**Equations 13-16**).

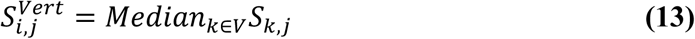

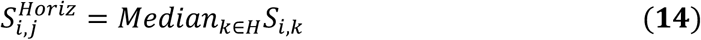

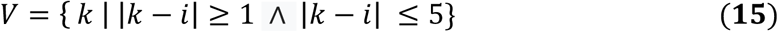

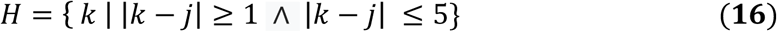

To compute the expected background signal for 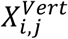, we multiplied the left and right footprint values by the bias factor and selected the maximum value (**Equation 17**). To compute the expected background signal for 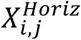, we multiplied the top and bottom footprint values by their bias factors and selected the maximum value (**Equation 18**).

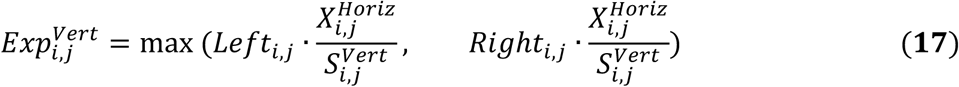

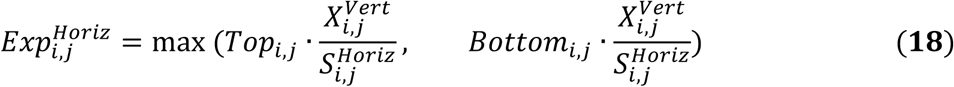

We assumed that the horizontal and vertical signals followed independent Poisson distributions parameterized by their background expected signals, testing whether the observed horizontal and vertical signals were higher than their expected signals (**Equations 19-20**).

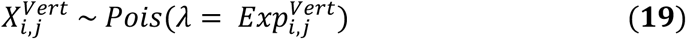

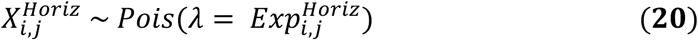

After computing p-values, we filtered and clustered significant pixels into stripes. First, we removed pixels with p-values greater than 0.15 and fold-changes less than 1.5. We also discarded pixels that had observed values less than 2 in the balanced contact matrix. After filtering, we clustered directly adjacent pixels in the vertical and horizontal directions separately. Horizontal and vertical clusters containing less than 3 pixels were thrown out. Then, we carried out a second clustering step where we merged surviving clusters with nearby clusters in the same direction. The maximum allowed distance between two clusters was 20 pixels. Once nearby clusters were merged, only those with 12 pixels or more were kept. Then, directly adjacent clusters in the orthogonal direction were merged together. The maximum cluster width was capped at 5 pixels. After clustering significant pixels in each time point, we filtered out clusters across time-points that were only called once. For a cluster to survive, it had to exist in at least two time-points. Any clusters that remained were considered to be real stripes.

#### Partitioning of stripe calls

Horizontal stripes and vertical stripes are analyzed separately. To identify horizontal stripes that contain inwardly oriented CTCF at their anchors, anchors were merged across all cell cycle stages to define the anchor region of a stripe. As long as the anchor region intersect with ≥ 1 CTCF peaks that contain down-stream oriented CTCF motif, we classify this stripe as one that contains inwardly oriented CTCF. All other horizontal stripes (no CTCF peak, oppositely oriented CTCF peak or CTCF peak without motif) are considered as stripes without inwardly oriented CTCF. Similar approach was applied to vertical stripe that target up-stream oriented CTCF peaks. To generate aggregate horizontal stripes, for the anchor region of each individual stripe, the central pixel (anchor region = odd number of pixels) or the upper one of the central two pixels (anchor region = even number of pixels) was identified. Then, a 10 * 200 pixel^2^ horizontal rectangle region was extracted from the 10kb binned, KR balanced, quantile normalized Hi-C contact map, originating from the central pixel. Average read counts of each pixel in the rectangle region was computed across all horizontal stripes as the aggregated horizontal stripe. Similar approach was taken to generate the aggregated vertical stripe. Aggregated horizontal and vertical stripes were then merged together to generate the aggregated stripe contact map.

### Loop calling

#### Expected modeling

We implemented an expected modeling strategy similar to that used by the HiCCUPS algorithm proposed by Rao et al. ^4^. For expected modeling and all subsequent loop calling steps, we restricted our analysis to bin-bin pairs with interaction distances of 10 Mb or lower, or equivalently, the first 1,000 diagonals of a 10 kb Knight-Ruiz balanced contact matrix. All computations related to expected modeling were performed on the balanced contact matrix *S*.

We first computed a one-dimensional distance-dependent expected model *D* by computing the average interaction count of each of these first 1,000 diagonals (**Equation 21**):

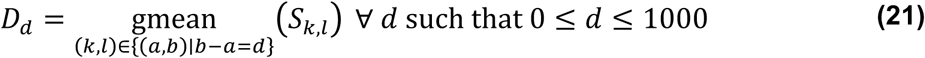

where *D*_*d*_ is the expected value for the interaction between two bins separated by *d* bins, *S* is the balanced contact matrix, and the geometric mean is computed over sets of bin-bin pairs separated by the same number of bins *b* – *a* = *d*. We compute 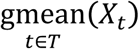 as the geometric mean over indices *t* taken from a set *T*, using a pseudocount of 1 to avoid the effects of zeros in the contact matrix (**Equation 22**):

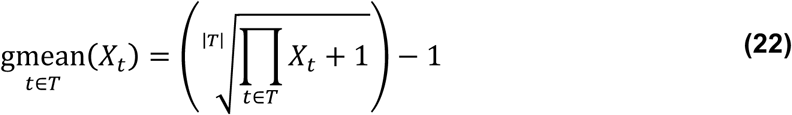

We next employed two distinct strategies for adjusting this one-dimensional distance-dependent expected matrix for local changes in contact frequency due to chromatin domain structure. For bin-bin pairs with interaction distances greater than 100 kb (beyond the first 10 diagonals of the contact matrix), we computed a correction factor using the donut and lower left filters employed by Rao et al. ^4^. The “footprints” of these filters represent local windows around each pixel at which we want to compute a corrected expected value. These windows specify a set of other nearby pixels which represent a good “local background” sample and will contribute to the correction factor. Mathematically, we can represent the footprints as sets of indices that identify which nearby pixels are included in the “local background” for that footprint. The donut footprint around bin-bin pair (*i,j*) is (**Equation 23**):

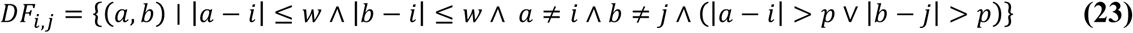

where *DF*_*i,j*_ represents the set of bin-bin pairs (*a, b*) that are included in the donut footprint centered on bin-bin pair (*i,j*), and *p* and *w* are parameters that control the inner and outer radius of the donut shape. Intuitively, bin-bin pairs (*a, b*) are included in the donut footprint around bin-bin pair (*i,j*) if they lie within a (2*w* + 1) × (2*w* + 1) square centered on (*i,j*) (first two conditions in the set construction in **Equation 23**), unless (i) they fall on the same row or same column as (*i,j*) (third and fourth conditions in the set construction in **Equation 23**) or (ii) they lie within a (2*p* + 1) × (2K*p*+ 1) square centered on (*i,j*) (fifth condition in the set construction in **Equation 23**). We used fixed constants *p* = 4 and *w*= 16 for this analysis.

We used the donut footprint to compute donut-corrected expected values as in Rao et al. ^4^ (**Equation 24**):

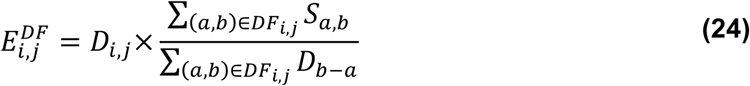

where *E*^*DF*^ represents the matrix of expected values corrected using the donut footprint. The fraction in this expression can be thought of as a correction factor that normalizes the one-dimensional distance-dependent expected value *D*_*b*–*a*_ to account for local enrichment or depletion of balanced values *S*_*a,b*_ in the local donut-shaped window relative to their own one-dimensional distance-dependent expected values *D*_*b* − *a*_ (where *b* − *a* represents the number of bins separating bin *a* and bin *b* the two bins whose balanced interaction value is quantified by *S*_*a,b*_, and *D*_*a*–*b*_ represents the expected interaction value for interactions between bins separated by this many bins). Points below the diagonal, beyond our maximum interaction distance of 10 Mb, or in low-mappability regions (as defined above) were excluded from contributing to any of the summations.

We also employed the lower left footprint as proposed by Rao et al., which contains only those points of the donut footprint that lie both below and to the left of each (*i,j*) th pixel (**Equation 25**):

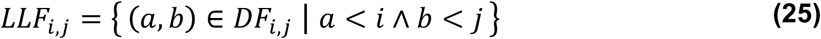

where *LLF*_*i,j*_ represents the set of bin-bin pairs (*a, b*) that are included in the lower left footprint centered on bin-bin pair (*i,j*). Note that to compute the lower left footprint *LLF*_*i,j*_, we start from the full donut footprint *DF*_*i,j*_ as described in (**Equation 23**) and keep only those points that lie below and to the left of the (*i,j*)th pixel. We used this lower left footprint to compute lower left corrected expected values as in Rao et al. (**Equation 26**):

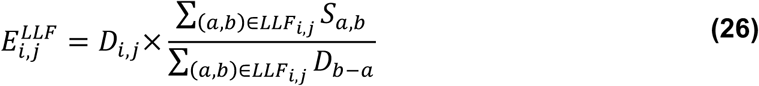

Finally, we computed final expected values for bin-bin pairs with interaction distances greater than 100 kb by taking the corrected expected value under whichever filter gave the higher expected value (**Equation 27**):

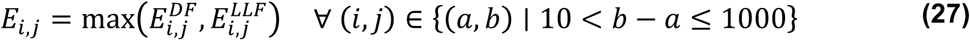

For bin-bin pairs with interaction distances within 100 kb, we observed that the normal donut filters were overestimating the expected value, reducing the sensitivity of loop calling near the diagonal of the contact matrix. Therefore, we used a different filter shape to model the expected values at these short interaction distances. Specifically, we used the upper triangle of the donut footprint, creating a new upper triangular footprint which only includes those bin-bin pairs (*a, b*)in the donut footprint that have interaction distances greater than or equal to the interaction distance of the entry for which the corrected expected value is being computed (**Equation 28**):

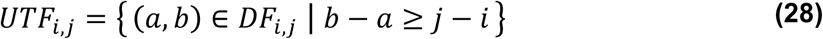

where *UTF*_*i,j*_ represents the set of bin-bin pairs (*a, b*) that are included in the upper triangular footprint centered on bin-bin pair(*i,j*).

We used this upper triangular footprint to compute upper triangular corrected expected values in analogy with the other footprints proposed by Rao et al. (**Equation 29**):

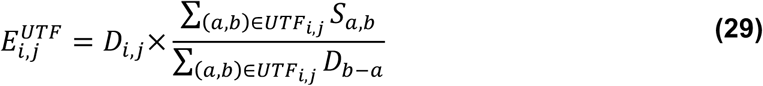

We used these upper triangular corrected expected values as our final expected values for all bin-bin pairs with interaction distances within 100 kb (**Equation 30**):

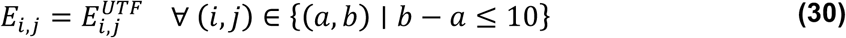

In summary, our matrix of final expected values *E*_*i,j*_ comes from the upper triangular corrected expected values for bin-bin pairs with interaction distances within 100 kb, and from the larger of the two other corrected expected values for bin-bin pairs with interaction distances beyond 100 kb (**Equation 31**):

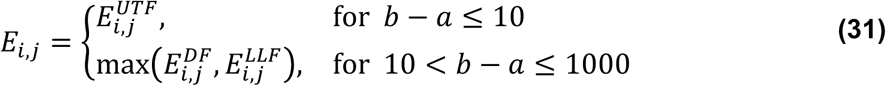

#### P-value calling

We called P-values for the raw read counts for each bin-bin pair under a Poisson statistical model, using the deconvolution approach proposed in Rao et al. ^4^. Briefly, we used the final expected value *E*_*i,j*_ and the Knight-Ruiz balancing bias vector ì to compute a biased expected value which is appropriate for comparison to the raw read counts x_*i,j*_ (**Equation 32**):

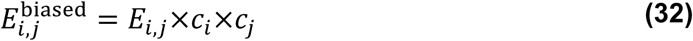

We then computed P-values against the null hypothesis that the raw read count *x*_*i,j*_ is less than or equal to the biased expected value 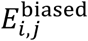 to assemble a matrix of P-values *p*_*i,j*_. Specifically, we compute the probability that the raw read count x_*i,j*_ is less than or equal to a Poisson-distributed random variable 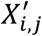 with mean 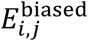 (**Equation 33**):

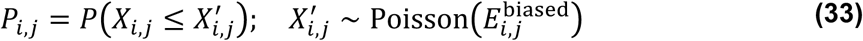

**Multiple testing correction.** We tested the null hypothesis in the previous section independently once for each bin-bin pair with interaction distance within 1 Mb. To perform multiple testing correction for this large number of tested hypotheses, we applied the lambda-chunking strategy proposed by Aiden and colleagues. Briefly, we first stratified bin-bin pairs (*i,j*)according to their biased expected values 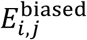 using logarithmically spaced bins with a bin spacing 2^1/3^. We then applied Benjamini-Hochberg false discovery rate control for the P-values *p*_*i,j*_ for each chunk separately to obtain a matrix of q-values *Q*_*i,j*_, which represent the maximum false discovery rate (FDR) at which an interaction would be called significant.

#### Clustering

The matrix of q-values *Q*_*i,j*_ computed in the previous section represents a quantitative measure of significance independently for each bin-bin pair. However, biological interactions often result in the elevation of contact frequency for several nearby bin-bin pairs. Therefore, we sought to identify clusters of nearby significant bin-bin pairs. First, we applied three initial thresholds to determine an initial set of significant bin-bin pairs. In order to be considered significant, a bin-bin pair (*i,j*)had to have a q-value *Q*_*i,j*_ ≤ 0.01 (corresponding to a false discovery rate of 1%), a balanced contact value *S*_*i,j*_ ≥ 10, and a fold-change between balanced contact value and final expected value 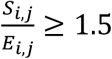. Next, we identified initial contact clusters, composed of directly-adjacent bin-bin pairs passing all three of our initial thresholds. To further reduce the possibility of false positives, we removed clusters with fewer than three significant bin-bin pairs.

We observed that this simple clustering strategy could sometimes include multiple visually-apparent loops in one large “supercluster”. In order to split these large clusters into smaller ones that are more likely to represent individual looping interactions, we applied progressively more stringent q-value thresholds (in order-of-magnitude steps from 0.01 to 1e-10 FDR) to our initial contact clusters. For each more stringent q-value threshold, we re-clustered the bin-bin pairs in the cluster that passed the new, more stringent q-value threshold. If there were no new clusters with at least three bin-bin pairs passing the more stringent q-value threshold, we stopped and kept the cluster as it was. If there was at least one new cluster with at least three bin-bin pairs passing the more stringent q-value threshold, we kept only those new, smaller clusters, and continued applying this rule recursively with progressively more stringent q-value thresholds. Thus, we refined our initial clusters into smaller clusters that represent sub-regions of relatively higher significance.

Finally, to further reduce the possibility of false positive interactions being called near the diagonal of the contact matrix, we removed all refined clusters containing a bin-bin pair whose interaction distance was within 20 kb. The final numbers of loops called at each time point is shown in (**Extended Data Fig. 6a**).

#### Merging loop calls across time points

We next sought to create a catalogue of interactions detected at any time point. We collected the set of bin-bin pairs (*i,j*) that were part of a final, refined cluster called in any time point, and clustered these into merged clusters of directly-adjacent bin-bin pairs. For each time point at which a merged cluster had any significant bin-bin pairs, we recorded the bin-bin pair (*i,j*) for which the log fold enrichment between balanced and expected values 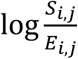 was the highest. We called the set of these most-enriched bin-bin pairs across all significant time points the “summit” of the loop. We used this summit to represent our best-guess of the exact epicenter of the loop, independent of time point.

### Downstream Analysis of Looping Interactions

#### Classification of merged loops using ChIP-seq and enhancer/promoter positions

We classified each loop according to the presence of ChIP-seq peaks, gene promoters, or enhancer marks at each of its two anchors. For the purposes of intersection with these marks, we defined the loop anchors as 20 kb regions centered on the respective midpoints of the two sides of the rectangle enclosing all the bin-bin pairs in the loop summit. If a loop had co-occupied CTCF and cohesin peaks under both anchors and that it does not simultaneously harbor cis-regulatory elements on both anchors, we placed it into the “structural loop” class. If a loop had putative enhancer marks or annotated promoters under both anchors, we placed it into the “E/P” class. If an E/P loop did not simultaneously harbor CTCF/cohesin co-occupied sites at both of its bases, then we term it as “E/P loop independent from CTCF/cohesin” otherwise, if an E/P loop had CTCF/cohesin co-occupying both of its bases, we term it as “E/P loop with CTCF/cohesin”. The “structural loop” class and the “E/P” class represent the two main classes of loops we used for our further analysis. Many of the remaining loops had CTCF peaks, putative enhancers, or annotated promoters at one or both anchors. Loops for each class are included in Extended Data Table 4.

#### K-means clustering of loops according to their looping dynamics

To quantitatively measure loop strength dynamics across time, we computed the average log fold enrichment 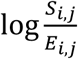 across all bin-bin pairs (*i,j*) in the summit of each loop at each time point. This allowed us to construct a matrix *A* for each loop class whose rows correspond to loops in that loop class, whose columns correspond to time points, and whose values *A*_*k,l*_ correspond to the strength (as quantified by average log fold enrichment) for loop *k* at time point *l*. We created a centered and scaled version of *A* by subtracting the row mean from each row and dividing each row by the row-wise standard deviation to obtain a new matrix *Ã* (**Equation 34**):

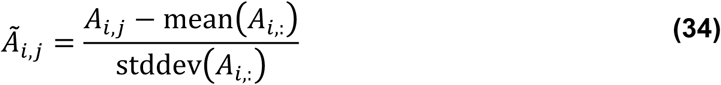

where *A*_*i*,:_ represents the 0th row of *A*. We chose this row-wise scaling and centering approach to highlight differences in temporal dynamics across loops. We then performed *k*-means clustering on the rows of *Ã*. We repeated this process for each of our two main loop classes. We observed that choosing *k* = 3 clusters recovered the most consistent and biologically interpretable clusters. We visualized the clustering by plotting a clustered heatmap of *Ã* (**Fig. 2h, 3e**).

#### Aggregate peak analysis (APA)

We performed aggregate peak analysis on each cluster at each time point for each loop class. Briefly, we extracted 130 kb x 130 kb squares from the quantile-normalized balanced contact matrices for each loop in each class and cluster at each time point. To avoid including the effects of the strong distance dependence background signal near the diagonal, we excluded loops with interaction distances less than 100 kb. We averaged the squares pixel-wise across loops within each class and cluster at each time point to obtain the APA plots in (**Extended Data Fig. 6e, h**). We also plotted APA plots for all loops in each loop class (“structural” and “E/P independent from CTCF/cohesin”) irrespective of their *k*-means clustering assignment, again making one plot for each time point while holding the set of included loops constant (**Fig. 3b**). To illustrate the progressive appearance of loops across time points, we performed a separate APA in which we grouped loops according to the time point at which the loop was first called significant. We plotted the average APA plot for each group of loops at each time point in (**Extended Data Fig. 6b**).

#### Unique anchor set construction

We sought to quantify the properties of loop anchors in different loop classes and *k*-means clusters, but loop anchors are often re-used by multiple loops (which may be in different loop classes or clusters). Therefore, we constructed sets of unique, non-redundant anchors that participated in loops from only one specific *k*-means cluster of the “E/P” loop class. First, we selected the final merged loops in the “E/P” loop class. Next, we separated each of these loops into its two constituent loop anchors, represented by 20 kb regions centered on the midpoints of the two sides of the rectangle enclosing all bin-bin pairs in the loop summit (matching the procedure used for the original loop classification above). We then merged loop anchors that overlapped by 10 kb or more, recording the original *k*-means clusters that the individual loop anchors belonged to. In a negligible number of highly complex looping regions, this merging process resulted in merged anchors larger than 40 kb in size; we discarded these merged anchors as too complex to resolve by our approach. Finally, for each *k*-means cluster, we selected the merged loop anchors used exclusively by loops from that *k*-means cluster to generate the final unique anchor set for that *k*-means cluster. This procedure is summarized in the schematic in (**Extended Data Fig. 8b**). The unique loop anchor sets for each *k*-means cluster were used to generate the histone modification ChIP-seq signal quantifications shown in (**Extended Data Fig. 8c**).

#### Capture-C data processing

Three biological replicates per probe were performed for each cell cycle stage (pro-meta to mid-G1). Raw reads were processed with publically available scripts^34,35,36^. Capture-C interactions were normalized to total interactions. Each biological replicate was individually analyzed and the resulting bigwig files were averaged together to create a final bigwig file for genome track visualization. Quantification of Capture-C read density at designated regions were performed on each biological replicate using the UCSC utility: BigWigAverageOverBed.

#### Promoter and enhancer annotation

We performed the following analysis to identify putative active promoters and enhancers in G1E-ER4 cells. H3K27ac raw ChIP-seq files on un-induced G1E-ER4 cells were acquired from GEO database (GSE61349) ^37^. Two H3K27ac replicates were pooled and peaks were called using MACS2 with parameter “-q 0.01” and further filtered for peaks with score >= 100. To this point we have 29602 peaks, which we partitioned into 9436 putative promoter peaks (<=1kb distance between peak center and TSS) and 20166 putative enhancer peaks (>1kb distance between peak center and TSS). We then stitched nearby promoter or enhancer peaks if the gap between them is less then 1kb, which resulted in 7456 promoter peaks and 14838 putative enhancer peaks. Additional external ChIP-seq data used in this study are listed as below: (1) H3K4me1 (GSM946535) ^38^, (2) H3K4me3 (GSM946533) ^38^, (3) H3K36me3 (GSM946529) ^38^, (4) H3K9me3 (GSM946542) ^38^ and (5) H3K27me3 (GSM946531) ^38^. Raw fastq reads associated with these of experiments were aligned to mm9 using Bowtie2, alignments with mapping quality QMap>10 were retained. PCR duplicates, mitochondria reads, reads mapped to contigs or ENCODE blacklist regions were removed using BEDtools. Bigwig files normlaized to RPKM were also generated by MACS2.

#### ChIP-seq alignment and peak calling

2-3 biological replicates of ChIP-seq experiments targeting CTCF or Rad21 were performed for each cell cycle stage and asynchronous populations. Input material corresponding to each time point or asynchronous populations were also sequenced. For reads alignment, fastq file per replicate per cell cycle stage was mapped to the reference mouse genome mm9 using bowtie (0.12.7). Reads were filtered to remove non-uniquely mapped reads and PCR duplicates. Bedgraph files in the unit of RPKM for each IP and corresponding Input libraries were generated by MACS2, and subtracted by MACS2 (“macs2 bdgcmp”). The resulting bedgraph files were converted to bigwig format using UCSC toolkit (“bedGraphToBigWig”). Negative values were replaced by zero. ChIP-seq signal of them in each replicate were summarized from replicate bigwig files using bwtool. Pearson correlation was then plotted among all CTCF replicates and Rad21 replicates respectively. The high correlation among biological replicates demonstrated the reproducibility of our ChIP-seq experiments, allowing us to merge reads from biological replicates. To do this, filtered reads were pooled together and down-sampled to equivalent read counts across all cell cycle stages. Peaks were identified using the MACS2 with punctate calling for both CTCF and Rad21 (p-values 1e-8 and 1e-4 respectively, Extended Data Table 5). To increase the confidence of identifying CTCF peaks in pro-metaphase population, we down-sampled filtered reads from 3 biological replicates of CTCF ChIP-seq on pro-metaphase samples to equivalent read counts and called peaks in each individual biological replicate with MACS (punctate calling with p-value 1e-6). Similar approach was applied to the two individual biological replicates of Rad21 ChIP-seq on pro-metaphase samples (diffuse calling, p-value 1e-6).

#### Generation and annotation of non-redundant peak list

After peak calling of CTCF and Rad21 on merged, down-sampled files at all cell cycle stages, a non-redundant union set of CTCF and Rad21 peak list was generated (Extended Data Table 6). The presence/absence of peaks in each merged sample were computed using BEDtools intersect. Peaks were annotated to closest genes using HOMER tools, and CTCF motif was scanned for in each union peak again using HOMER tools. For the entire union set of peaks and selected subsets of peaks, CTCF or Rad21 ChIP-seq signal were summarized over them using bwtool. Mitotically retained CTCF was identified as long as a CTCF peak was called in at least two of the three biological replicates at pro-metaphase. Mitotic Rad21 peaks was defined as long as a Rad21 peak was called in both biological replicates during pro-metaphase. For motif enrichment analysis of M-and IO-sites, peak centers of the 7741 M-and 33958 IO-sites were identified and expanded to -250bp to +250bp around the center. DNA sequence of these regions were acquired with BEDtools-getfastq and used as input to feed into MEME-ChIP ^39^, for combinatorial analyses of de novo motif enrichment and comparisons of motif with known motif database.

#### Recovery dynamics of CTCF/cohesin co-occupying bin-bin pairs

We implemented a previously established pipeline to unbiasedly intersect ChIP-seq with Hi-C datasets ^40^. To examine the impact of both CTCF and Rad21 on chromatin interaction, we took the 10kb binned, Knight-Ruiz balanced, quantile normalized contact map from all cell cycle stages and intersected with a list of 19520 binding sites co-occupied by CTCF and Rad21. 10kb interacting bin-bin pairs were extracted as long as both bins contain CTCF/Rad21 co-occupied binding sites. We filtered the 10kb interacting bin-pairs based on (1). the observed over expected values of late-G1 must be > 2 to ensure that we were analyzing solid interactions and (2). the difference between late-G1 and pro-meta (obs/exp(late-G1)-obs/exp(pro-meta)) must be greater than 0.1. We then, constructed a recovery curve for each interacting bin-bin pair. Briefly, the observed over expected values of each cell cycle stage was subtracted by that of pro-metaphase (obs/exp(any time point)-obs/exp(pro-meta)) and then divided by the increment from pro-meta to late-G1 (obs/exp(late-G1)-obs/exp(pro-meta)). By doing this, the recovery curve would start from 0 at pro-metaphase and reach to 1 at late-G1 phase. Individual pin-bin pairs were then parsed into 10 groups based on their genomic separation (10-100kb, 110-200kb …… 910-1000kb). The recovery rate of interacting bin-bin pairs within the same distance group were then averaged together to create the overall recovery rate of that distance range.

### Data Availability

Hi-C and ChIP-seq raw and processed data will be deposited to GEO database for public access.

**Extended Data Fig. 1.**
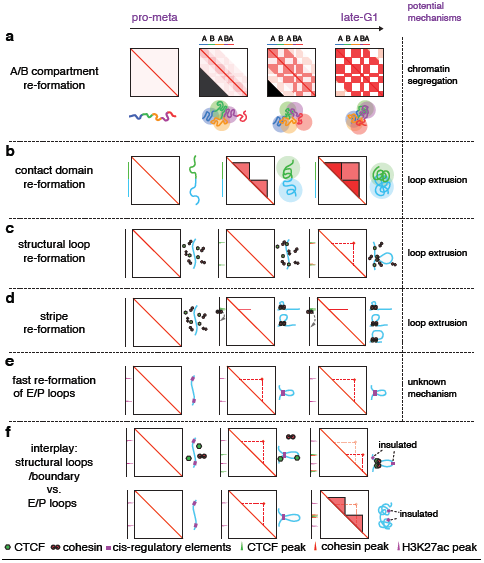
Models of post-mitotic chromatin re-organization. **a**, Schematic illustration of the gradual emergence, intensification and expansion A/B compartments (checkerboards) from pro-M to late-G1 phase, coupled with schematics of chromatin topology. **b**, Sub-TADs (smaller triangles) emerge first after mitotic exit, followed by converging into a bigger TAD (big triangle). **c**, Formation of a structural loop coincides with the positioning of cohesin, but not CTCF after mitosis. **d**, The gradual extrusion of cohesin complex along DNA fiber from one anchor point with CTCF, reflected as enrichment of interactions between the anchor and a continuum of DNA loci on the contact map. **e**, Fast post-mitotic enrichment of interactions between two DNA loci, demarcated by cis-regulatory elements. **f**, The interplay between transiently “mis-wired” cis-regulatory contacts and structural loops. In the model, cis-regulatory elements rapidly form contacts after mitosis, which is thereafter insulated by the formation of an inner domain boundary or interfered by a nearby structural loop.

**Extended Data Fig. 2.**
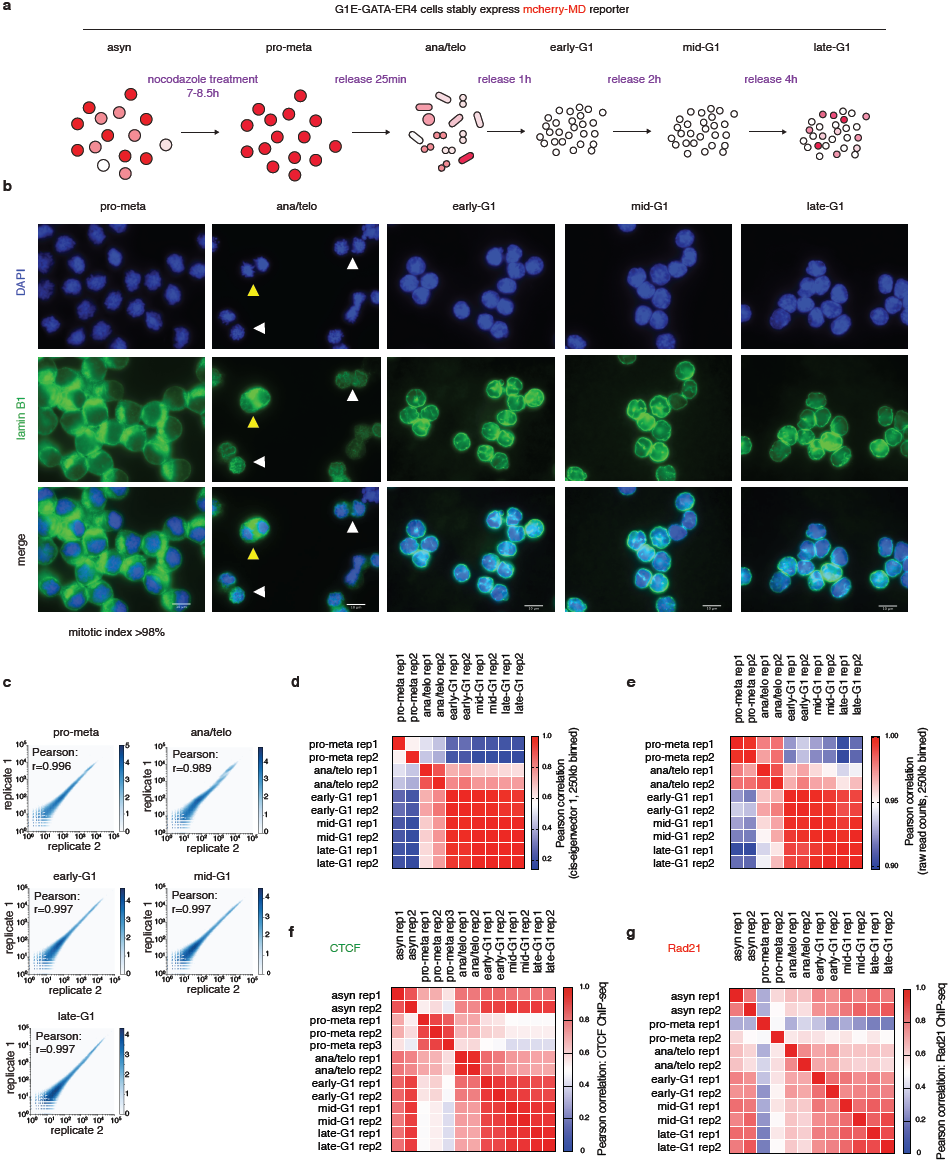
Experimental workflow and immunofluorescent staining of purified cells and HiC and ChIP-seq quality control. **a**, A schematic showing expected mcherry signal change during nocodazole synchronization and release. **b**, Representative images showing DAPI and lamin B1 staining of FACS purified cells across all cell cycle stages. Nuclear envelop (demarcated by lamin B1 staining) was disassembled in pro-metaphase and re-assembled in ana/telophase. Note that mitotic index of pro-metaphase cells after FACS purification is >98%. Yellow and white triangles indicate anaphase and telophase cells respectively. Scale bar: 10μm. **c**, Hexbin plots showing the high correlation of Hi-C data between two biological replicates across all cell cycle stages. Bin size: 250kb. **d**, A heatmap showing the Pearson correlation coefficient among all samples based on the PC1 value of 250kb bins. **e**, A heatmap showing the Pearson correlation coefficient among all samples based on raw read counts. Bin size: 250kb. (**f**-**g**), Two heatmaps, showing Pearson correlation coefficient of CTCF and Rad21 ChIP-seq among all samples, respectively.

**Extended Data Fig. 3.**
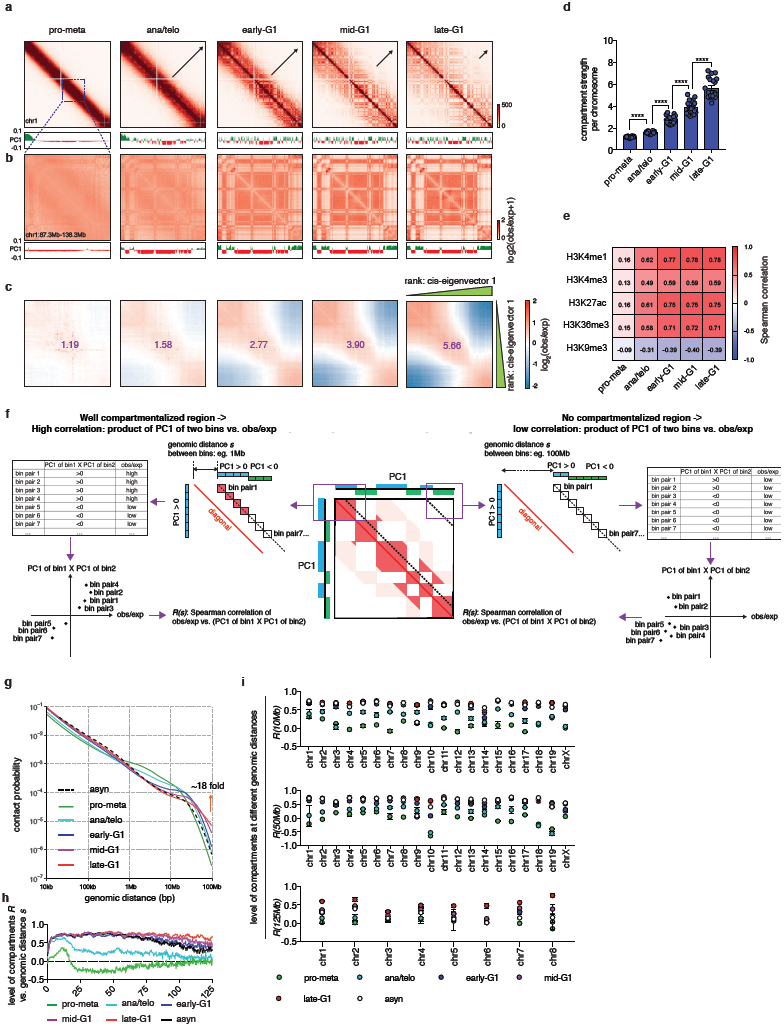
Early detection and progressive intensification and expansion of A/B compartment formation after mitosis. **a**, KR balanced Hi-C contact maps, showing the restoration of chromatin A/B compartments in chromosome 1 after mitosis, along with genome browser tracks showing cis-eigenvector 1 values across all cell cycle stages. Bin size: 250kb. **b**, A zoomed in region (chr1:87.3M-138.3M) of (**a**) to show the clear plaid like compartment pattern in ana/telophase, which is further intensified over time. PC1 track illustrates the early detection of A/B compartments by cis-eigenvector decomposition. **c**, Saddle plots showing the progressive gain of compartment strength over time. The heatmaps show chromosome averaged distance normalized interaction frequency between pairs of 250kb bins in *cis*, ranked in ascending order of the cis-eigenvector 1 values of late-G1 phase samples on x and y axis respectively. The upper left corner of each heatmap shows the interactions between pairs of bins that both belong to B-type compartment. The lower right corner of each heatmap shows the interactions between pairs of bins that both belong to A-type compartment. Upper right and lower left corners of each heatmap show interactions between pairs of bins that separately belong to different types of compartments. The purple numbers on each heatmap denotes compartment strength of that cell cycle stage. **d**, Barplot showing the compartment strength of each individual chromosome over time. **** *p*<0.0001 Wilcoxon test. **e**, A heatmap, showing the Spearman correlation coefficients of genome wide cis-eigenvector 1 values and indicated histone mark signals. **f**, A schematic showing how levels of compartmentalization *R(s)* are calculated at different distance scales (eg. 1Mb or 100Mb). Each dotted line indicates a series of 250kb bin-bin pairs that are separated by a given genomic distance *s* (the distance from the diagonal to the dotted line) within a chromosome. For all bin-bin pairs separated by distance of *s*, a Spearman correlation coefficient *R(s)* was generated between obs/exp and the product of two cis-eigenvector 1 values (PC1 (bin1) X PC1 (bin2)). *R(s)* is expected to be high in well compartmentalized regions (left panel) and low at long distances with no compartments (right panel). **g**, Chromosome averaged distance dependent contact frequency *P(s)* plots for all cell cycle stages plus asynchronous sample. Orange arrow demarcates the trend of contact frequency change at super long (∼100Mb) genomic distance during pro-M-G1 transition. **h**, *R(s)* decay curve of chromosome 1 across all cell cycle stages. In ana/telophase, *R(s)* display rapid decay as *s* increases beyond ∼12Mb, suggesting restricted compartmentalization. After G1 entry, such decay is significantly attenuated. In late-G1 phase, *R(s)* curve still maintain constant when *s* increases beyond 100Mb, suggesting extensive expansion of the plaid like compartment pattern. **i**, *R(s)* of each individual chromosome across all cell cycle stages when *s* equals to 10, 50 and 125Mb. Error bar denotes SEM from two biological replicates of Hi-C experiments.

**Extended Data Fig. 4.**
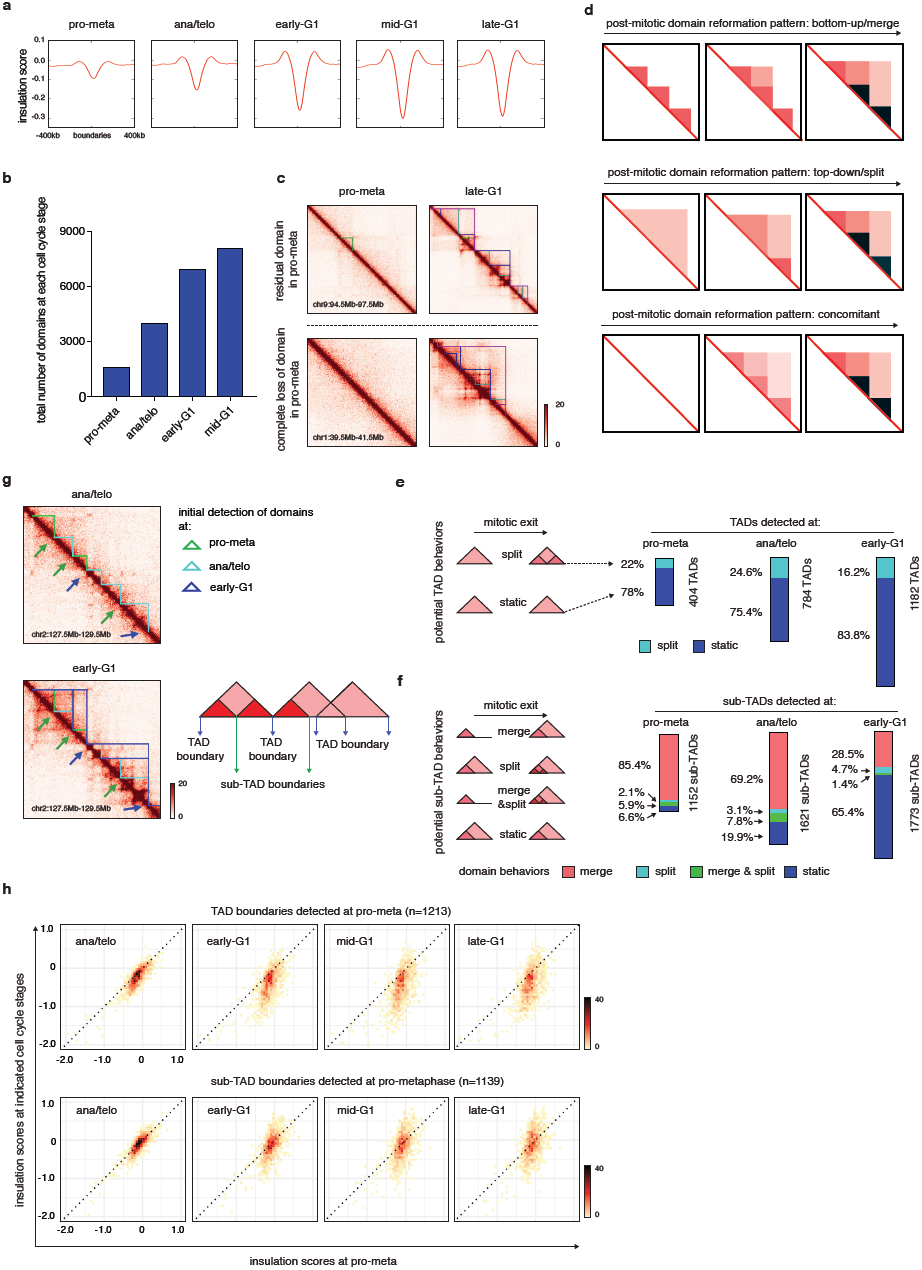
Behaviors of TADs and sub-TADs after mitosis. **a**, meta-region plots of insulation scores centered around domain boundaries across all cell cycle stages. **b**, Barplot showing the number of domain calls from pro-metaphase to mid-G1 phase. **c**, Upper panel: KR balanced, quantile normalized Hi-C interaction maps showing residual domain like structural in pro-metaphase (indicated by green lines). Lower panel: Hi-C interaction maps show a representative region of complete disruption of domains in pro-metaphase. Bin size: 10kb. **d**, Schematics showing the possible patterns of domain hierarchy reformation: bottom-up/merge, top-down/split and concomitant. **e**, Left panel: A schematic illustration of the two possible behaviors of TADs after emergence: static and split. Right panel, Barplots showing the fraction of TADs that display either type of behavior after detection at pro-metaphase, ana/telophase and early-G1. **f**, Left panel: A schematic illustration of the four possible behaviors of sub-TADs: merge, split, merge & split and static. Right panel: Barplots showing the fraction of sub-TADs that display each type of behavior after detected at pro-metaphase, ana/telophase or early-G1. **g**, Left panel: KR balanced, quantile normalized Hi-C contact map showing the insulation change of representative TAD-and sub-TAD-boundaries from ana/telophase to early-G1. Sub-TAD-and TAD-boundaries are indicated by green and blue arrows respectively. Bin size: 10kb. Right panel: A schematic showing how TAD-and sub-TAD-boundaries are defined. **h**, Bin plots showing the insulation score change of 1213 TAD-boundaries (upper panel) and 1139 sub-TAD-boundaries (lower panel) that are detected at pro-metaphase. Pro-metaphase insulation scores are plotted on x axis, against insulation scores of ana/telophase, early-, mid-and late-G1 respectively.

**Extended Data Fig. 5.**
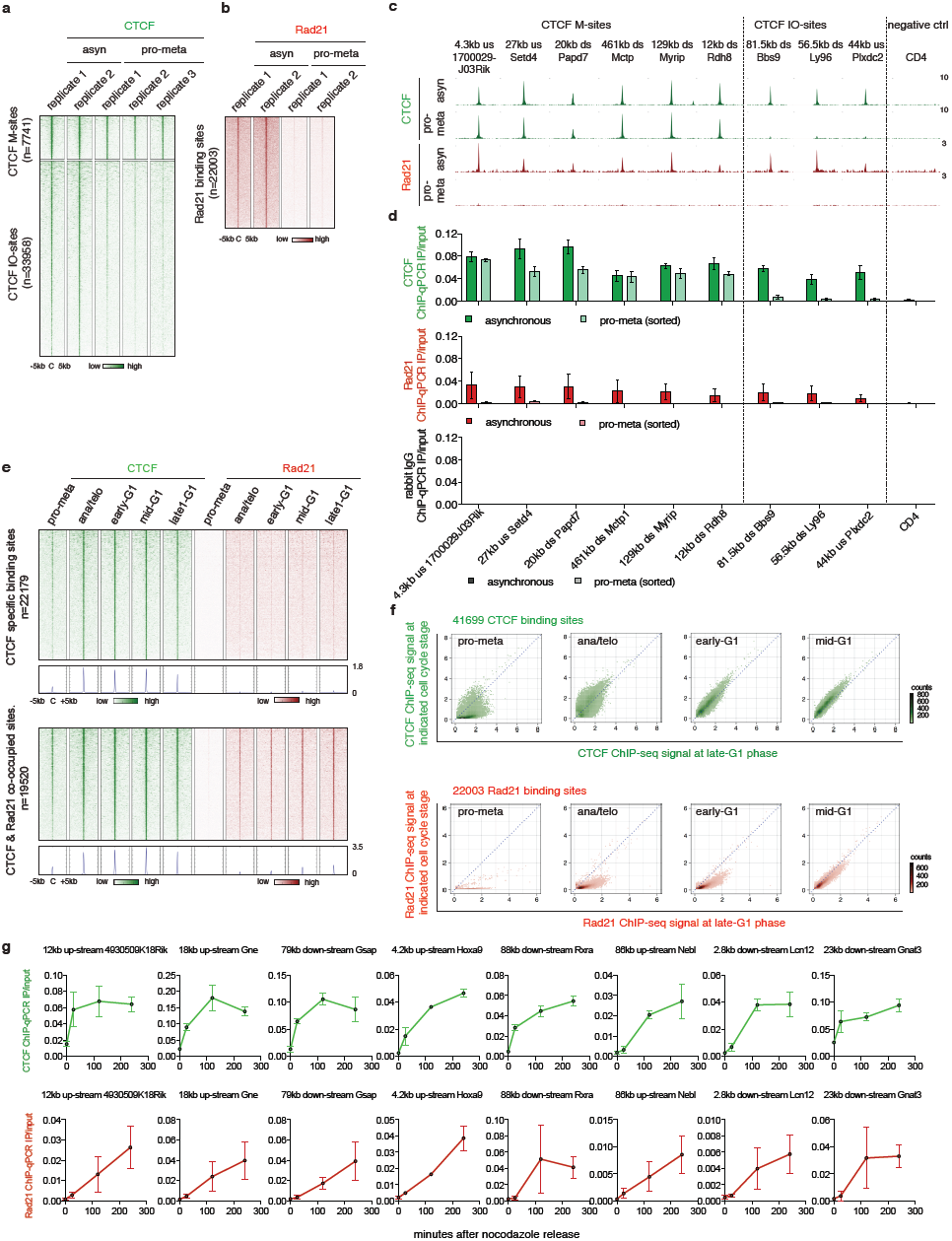
CTCF displays mitotic retention and fast recovery dynamics than cohesin after mitosis. **a**, A density heatmap of CTCF ChIP-seq from all biological replicates of asynchronous and pro-metaphase samples, centered around all CTCF peaks subdivided into M-and IO-CTCF binding sites. **b**, A density heatmap of Rad21 ChIP-seq from all biological replicates of asynchronous and pro-metaphase samples centered around all Rad21 peaks. **c**, Genome browser tracks showing CTCF and Rad21 ChIP-seq signals of asynchronous and pro-metaphase samples at indicated regions. **d**, ChIP-qPCR experiments of CTCF, Rad21 and rabbit IgG control in asynchronous and pro-metaphase samples as independent validation of CTCF mitotic retention and loss of Rad21 specific binding in pro-metaphase. Error bars denote SEM from 2-3 biological replicates. **e**, A density heatmap and metagene plots of replicate averaged CTCF and Rad21 ChIP-seq across all time points centered around CTCF specific and CTCF/Rad21 co-occupied binding sites. **f**, Bin plots showing the replicate averaged ChIP-seq signals of CTCF and Rad21 peaks for each cell cycle stage (y-axis) against late-G1 (x-axis). **g**, ChIP-qPCR experiments of CTCF and Rad21 at indicated loci in pro-metaphase, ana/telophase, mid-and late-G1 samples as independent validation of the differential post-mitotic re-positioning dynamics between CTCF and cohesin.

**Extended Data Fig. 6.**
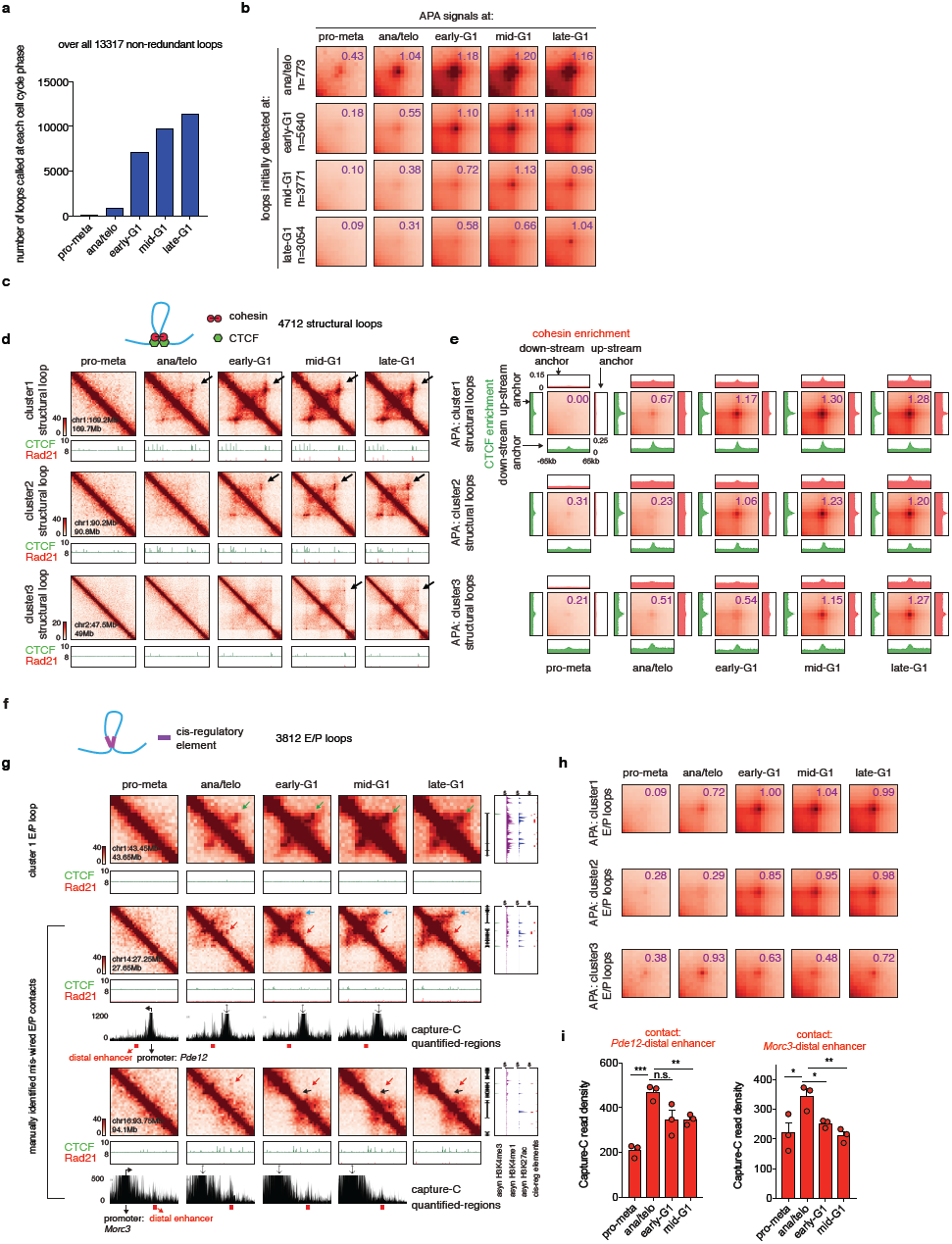
Loop statistics and k-means clustering on structural loops and E/P loops. **a**, Barplot showing the number of loop calls at each cell cycle stage. **b**, Aggregated peak analysis (APA) of loops that are newly detected at each cell cycle stage (from ana/telo to late-G1). Purple numbers on each heatmap denote average loop strength. Bin size: 10kb. **c**, A schematic of structural loops. Both anchors of structural loops are co-occupied by CTCF and cohesin. In addition, structural loops do not possess cis-regulatory elements on both anchors simultaneously. **d**, KR balanced, quantile normalized Hi-C interaction maps showing representative regions that contain cluster 1 (chr1:169.2M-169.7M), 2 (chr1:90.2M-90.8M) and 3 (chr2:47.5M-49M) structural loops respectively. Bin size: 10kb. Loops are indicated by black arrows. Contact maps are coupled with genome browser tracks showing CTCF, cohesin across all cell cycle stages. **e**, APA of cluster1, 2 and 3 structural loops across all cell cycle stages. Each APA heatmap is coupled with four metagene plots corresponding to piled up CTCF and Rad21 ChIP-seq signals centering around either up-stream or down-stream loop bases. Bin size: 10kb. Purple numbers indicate average loop strength. **f**, A schematic of E/P loops. **g**, KR balanced, quantile normalized Hi-C interaction maps showing an additional example of cluster 1 (chr2:43.45Mb-43.64Mb, loop indicated by green arrow) and two manually identified examples of transiently mis-wired E/P contacts (*Pde12* locus and *Morc3* locus, indicated by red arrow). Bin size: 10kb. Heatmaps are coupled with genome browser tracks of CTCF and cohesin across all time points as well as asynchronous H3K4me3, H3K4me1 and H3K27ac and annotations of cis-regulatory elements. Boundaries or structural loops anchors that interfere with mis-wired E-P contacts are indicated by black and blue arrows respectively. Capture-C interaction profiles of *Pde12* and *Morc3* loci are also shown. Probes (anchor symbol) are located at promoters of *Pde12* and *Morc3* genes respectively. Red bars demarcate the interaction between promoters and distal enhancers. **h**, APA of cluster 1, 2 and 3 (mis-wired) E/P loops across all cell cycle stages. Purple numbers indicate average loop strength. **i**, quantification of the Capture-C read density of the red regions in (**g**). Error bars denote SEM from three independent biological replicates. * *p*<0.05, ** *p*<0.01, *** *p*<0.001, Student’s t test.

**Extended Data Fig. 7.**
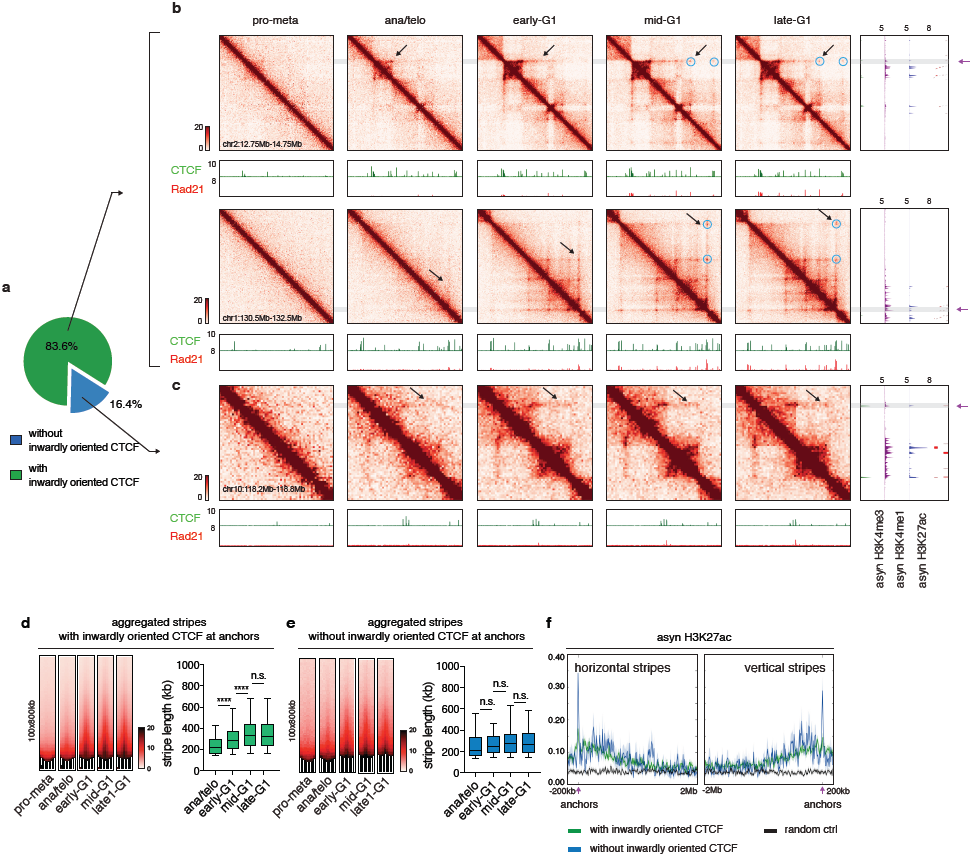
Reformation of chromatin stripes after mitosis. **a**, Pie chart showing the fraction of stripes with inwardly oriented CTCF at stripe anchors. **b**, KR balanced, quantile normalized Hi-C interaction maps showing two representative regions (chr2:12.75Mb-14.75Mb, chr1:130.5Mb-132.5Mb) that contain stripes with inwardly oriented CTCF at anchors. Bin size: 10kb. Contact maps are coupled with genome browsers tracks of CTCF and Rad21 across all cell cycle stages and tracks of asynchronous H3K4me3, H3K4me1 and H3K27ac and annotation of cis-regulatory elements. The dynamic extruding of stripes is indicated by black arrows. Stripe anchors are indicated by purple arrows. Punctuated loops along and at far end of stripes are indicated by blue circles. **c**, similar to (**b**) Hi-C interaction maps showing a representative stripe (chr10:118.2Mb-118.6Mb) that does not have inwardly oriented CTCF at stripe anchor. **d**, Left panel: aggregated Hi-C contact maps that compiles all stripes with inwardly oriented CTCF to show the overall dynamic growing of these stripes after mitosis. Right panel: boxplots showing the lengths of stripes detected at indicated cell cycle stage. 5 to 95 percentile were plotted. **** p<0.0001 Mann-Whitney test. **e**, Left panel: aggregated Hi-C contact maps that compiles all stripes without inwardly oriented CTCF. Right panel: boxplots showing the length these stripes detected at indicated cell cycle stage. 5 to 95 percentile were plotted. **f**, Asynchronous H3K27ac ChIP-seq file is plotted -200kb to 2Mb around the horizontal stripe anchors and -2Mb to 200kb around the vertical stripe anchors. Anchor position is indicated by purple arrows.

**Extended Data Fig. 8.**
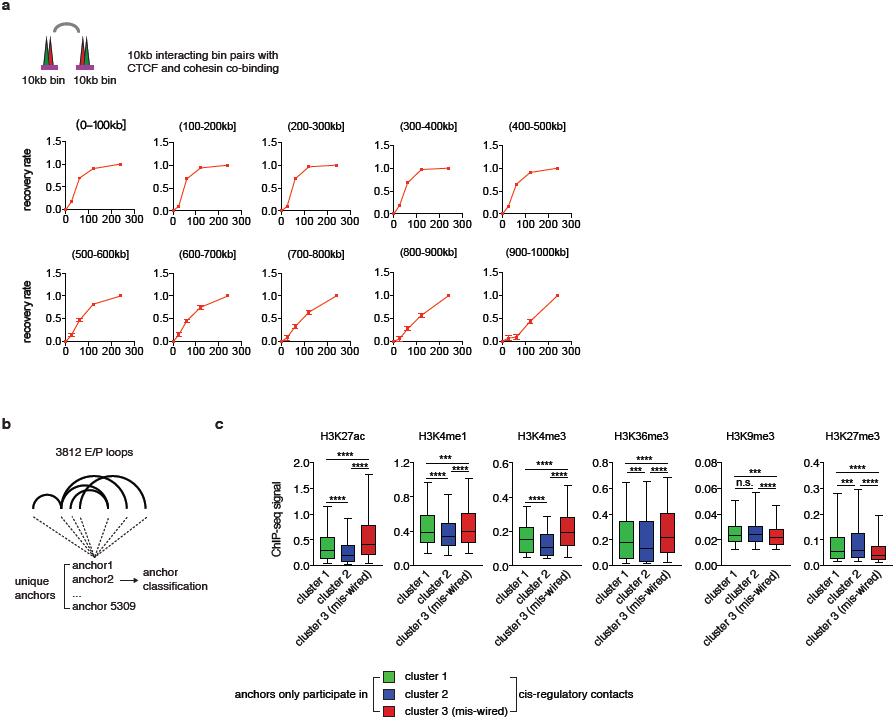
Additional evidence of CTCF/cohesin dependent and independent mechanisms underlying chromatin contact formation. **a**, Genome averaged recovery rate of, distance normalized interaction frequencies. Each individual plot represents aggregation of 10kb interacting bin-bin pairs that are separated by indicated genomic distance intervals. **b**, A schematic showing classification of anchors of E-P loops. **c**, Boxplots showing ChIP-seq signals of indicated histone modifications at anchors that solely participate in cluster 1, 2 or 3 (mis-wired) cis-regulatory contacts. 5 to 95 percentile were plotted. *** *p*<0.001, **** *p*<0.0001, Mann-Whitney test.

**Extended Data Table. 1.** Oligo sequences.

**Extended Data Table. 2.** Hi-C data processing statistics.

**Extended Data Table. 3.** Overall domain calls and domain calls at each cell cycle stage.

**Extended Data Table. 4.** Overall loop calls.

**Extended Data Table. 5.** CTCF & Rad21 ChIP-seq data processing statistics.

**Extended Data Table. 6.** CTCF & Rad21 union peak list.

